# Predictors of extinction risk in large tropical forest mammals: from global to local

**DOI:** 10.1101/2025.08.21.669653

**Authors:** Simon D. Schowanek, Douglas Sheil, Lydia Beaudrot, Pierre Dupont, Santiago Espinosa, Vittoria Estienne, Julia E. Fa, Jonas Geldmann, Patrick A. Jansen, Steig E. Johnson, Francesco Rovero, Fernanda Santos, Asunción Semper-Pascual, Andrea Fernanda Vallejo Vargas, Jorge A. Ahumada, Emmanuel Akampurira, Rajan Amin, Robert Bitariho, Adeline Fayolle, Davy Fonteyn, Ilaria Greco, Marcela Guimarães Moreira Lima, Matthew Scott Luskin, David Kenfack, Emanuel H. Martin, Eustrate Uzabaho, Cédric Vermeulen, Richard Bischof

## Abstract

Studies can only guide conservation if their findings are informative at the scales at which practitioners and policymakers operate. Yet, it is rarely tested whether large-scale studies reach similar conclusions to the smaller-scale studies on which conservation traditionally relies. We examine whether predictors of extinction risk are consistent across global, regional, and local scales, for 210 tropical forest mammal species (≥ 1 kg) that existed during the last 130,000 years, in 64 tropical forests, across three biogeographical realms. We found consistent predictors of extinction risk (body mass, generation length, diet, brain volume and scansoriality) when analyses differed only in their spatial resolution. However, predictors differed when analyses also varied in their temporal extent. Macroecological findings about extinction risk can, thus, inform conservation at smaller scales, but they risk misidentifying threatened species if differences in temporal extent are not recognised.

## Background

Humans are causing species extinctions at a rate unprecedented in recorded history [1,2]. Conservationists aim to mitigate these extinctions through practice (conducting actions on the ground), policy (committing to a course of action), and research (generating knowledge to guide practice and policy). While practice and policy can happen at various spatial or temporal scales, they are primarily implemented at national, sub-national, or local scales (e.g. inside country borders or protected areas), and over comparatively short timescales (e.g. months, years). In contrast, scientific studies that can inform conservation are conducted across various scales. While some studies use spatio-temporal scales similar to those encountered by policy makers and practitioners, others use larger scales, for example global or continental spatial scales or centennial to millennial timescales [3–6]. Due to this scale mismatch, scientific research at large spatio-temporal scales is often perceived as irrelevant for practical conservation [3,5–7].

To ensure large-scale studies become relevant for conservation, their findings must be informative at the scales at which practitioners and policymakers act [3,4]. This requires that the patterns described by scientific studies are scale generalisable (i.e. findings from one scale can be applied at another [8]) or alternatively that we understand why patterns vary across scales. For example, if palaeoecological studies show that, globally, larger-bodied species have been more vulnerable to extinction over millennial timescales [9], scale generalisation would suggest that body size has similar effects in present-day situations encountered by conservationists (e.g, see Rovero et al. [10] who describe a loss of large-bodied mammals close to human activities inside several present-day protected tropical forests). Yet, while scale generalisation (or the absence thereof) is often assumed, such assumptions are rarely tested.

Comparing findings across scales is challenging because trade-offs often exist between study resolution or grain (the smallest unit of observation) and extent (the spatial or temporal range covered [8]). Small-scale studies offer high resolution but lack the spatial or temporal coverage to permit more general conclusions. Large-scale studies, on the other hand, cover broader extents but typically lack the resolution needed to describe patterns in detail, which are needed for implementing local conservation practices. However, advances in automated monitoring methods, like camera trapping, increasingly offer datasets with extensive geographical coverage and high data resolution [11]. Although such datasets remain biased towards sites of high conservation interest, such as protected areas [12], they can provide extensive and detailed views of ecological patterns that may help bridge the gap between research at large scales and application at smaller scales.

Predicting species’ extinction risk at different scales offers an excellent means to test the generalisability of findings across these scales. First, preventing extinction is at the heart of conservation and studies of extinction are conducted on several spatio-temporal scales, ranging from global (e.g. [13]), to regional (e.g.[14]) to local (e.g. [15]). However, some have questioned the usefulness of large-scale studies of extinction risk, arguing that the findings of such studies are often inconsistent and have led to few clear and actionable conservation messages [16]. Conversely, local, and short-term studies are more easily affected by shifting baselines [17], which can obscure ecological and evolutionary patterns [18,19].

Second, extinction is a useful study system because, in theory, species losses at different scales are causally connected. A species’ extinction at large scales results from local extinctions throughout its range, suggesting that the patterns and predictors of extinction should, to some degree, be comparable across scales. Nonetheless, several social, demographic, and ecological patterns are scale-variant, meaning that the same phenomenon can effectively be treated as independent phenomena when analysed at different scales [8]. For example, while a species’ global range comprises the space use of all individuals, the space use of any given individual may teach us little about a species’ global range, and vice versa. Understanding which patterns are scale-variant and which are not will help improve the practical utility of large-scale studies in informing local conservation.

Here, we test whether extrinsic and intrinsic predictors of extinction risk in medium to large (≥ 1 kg) tropical forest mammals are generalisable across global, regional, and local scales. Using a Bayesian framework, we model the survival probability (the inverse of extinction probability) of 210 species in 64 tropical rainforest sites (protected: n = 31; partially protected: n = 25; unprotected, n = 8) across three biogeographical realms at these three scales (Fig. 1). The model consists of three logistic regressions, representing studies of survival at the different scales. The global regression describes which species survived across the tropics since the Eemian period, which started 130,000 years BP, providing a macroecological comparison with limited human impacts. The regional regression describes which species have persisted within a region. Regions are defined as similarly sized areas that broadly share a biogeographical and administrative history (see Methods). Most regions correspond to countries and resemble the administrative entities with which national and international conservation policy is concerned. The local regression describes which species have persisted at a given study site (e.g. a protected area).

**Figure 1.**
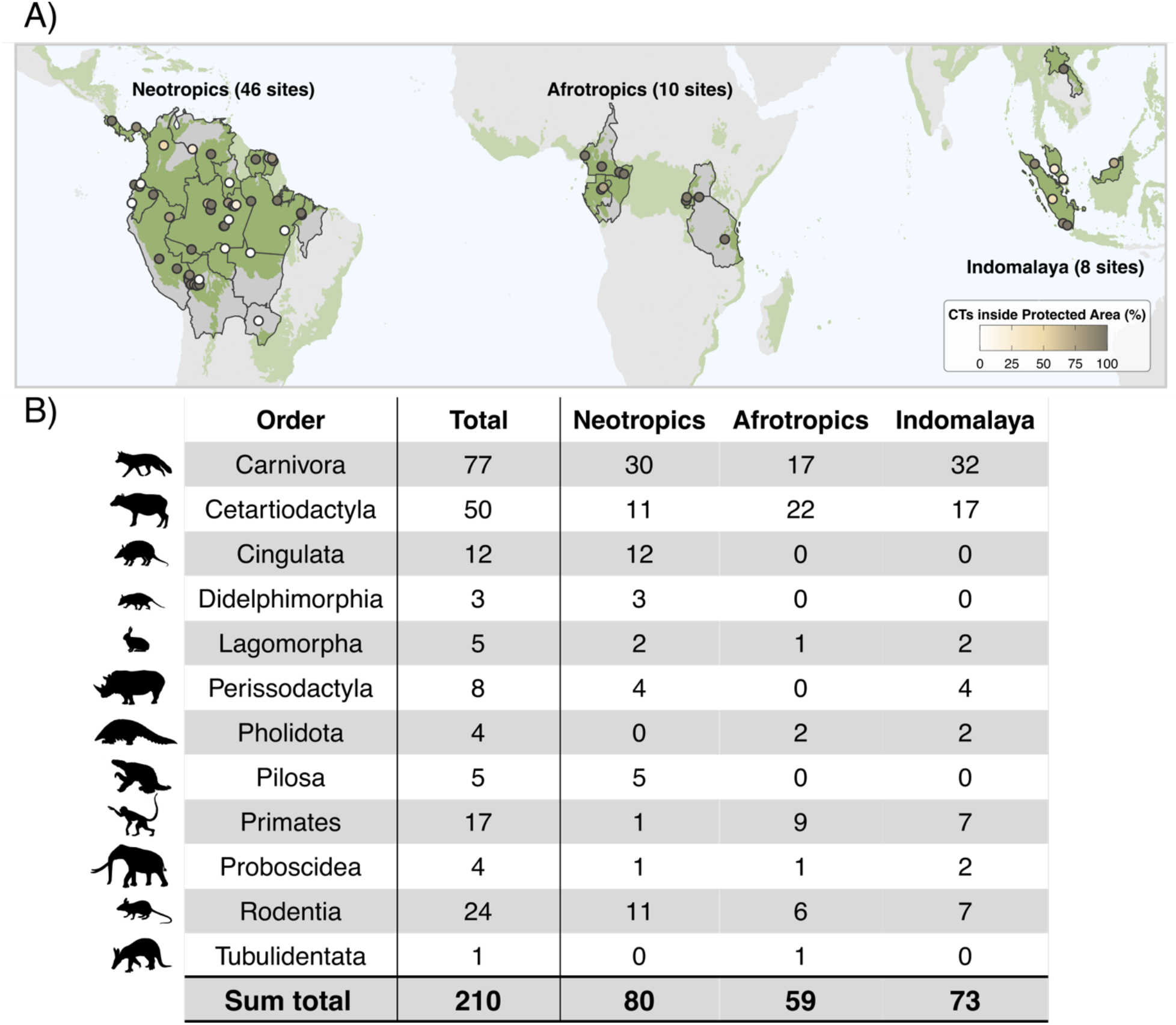
Biogeographic realms, regions, and sites and the tropical forest mammals studied. (**A**) The locations of the 64 sites used in this study, as well as the regions in which they are located (highlighted areas). Regions are approximately evenly-sized areas that broadly share a biogeographical history, and fall under the same administrative entities (see Methods). The color of the points shows how many of the sites’ camera traps are located inside a protected area. The green layer shows the distribution of tropical forest. **(B)** A summary of the species used in this study, summarized by order and biogeographical realm. Note that some species occur in multiple realms.

To compare how extinction risk predictors vary over space and time, we run two versions of this model (Fig. 2). In the spatial-only model, the only difference between regressions is their spatial resolution. The species pool is the same at each scale, though fewer species survive at smaller scales. In the spatio-temporal model, the spatial resolution changes as well as the temporal extent. We do so by excluding species from the species pool at smaller scales if they have already gone extinct at a higher scale. This gives each regression different but nested species pools. Excluding these species broadly corresponds to the chronology of mammal extinctions and the subsequent absence of species from scientific studies due to shifting baselines [17]. In the spatio-temporal model, the global, regional and local scales, therefore, correspond to studies with long, intermediate and short temporal extents, respectively. In both models, the regressions have the same pantropical extent.

**Figure 2.**
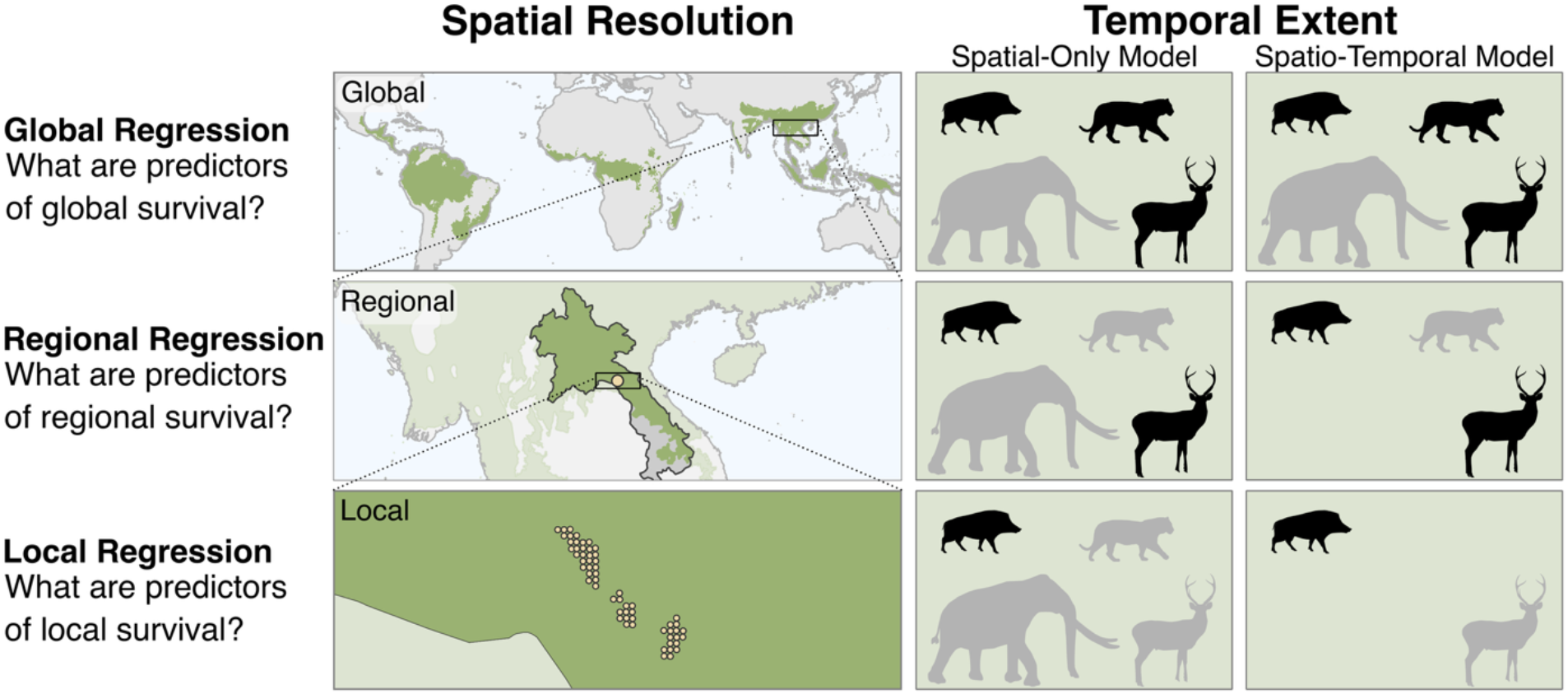
Framework used to test the similarity of predictors of species’ extinction risk across spatial and temporal scales. Column one describes the three regressions used in each model. Column two shows the spatial resolution of each regression. Columns three and four represent the species pools at different scales for the spatial-only and the spatio-temporal models, respectively. Black silhouettes represent extant species. Gray silhouettes represent extinct species. The spatial-only model changes the spatial resolution from global to regional to local but the temporal extent remains identical for each regression. In contrast, the spatio-temporal model changes the spatial resolution as well as the temporal extent. lt sequentially removes extinct species resulting in nested species pools for each regression.

We determined species’ presences and absences by combining present-natural ranges from PHYLACINE v1.2.1 (estimates of where species would live in the absence of human impacts[20]), ranges from the International Union for Conservation of Nature (IUCN), and camera trap observations (see Methods). We included three types of predictor variables: biogeographical variables (biogeographical realm), species traits (body mass [20], relative brain volume [21,22], relative generation length [23], vertebrate carnivory [20], and scansoriality [24,25]), and site variables (forest cover [26], human population density [27], protected area coverage [28]). We included site variables in the local regression only to account for their known effects on extinction risk rather than to test their consistency across scales. We also ran an alternative version of both models, which included phylogenetic variables, accounting for relatedness between species (See Methods). We used a reversible-jump Markov Chain Monte Carlo approach to quantify the probability that a predictor should be included in a regression [29]. The resulting inclusion probability provides a metric of the evidential support for each predictor (see Methods). We considered variables significant if they were more likely than not to be included in a regression (inclusion probability > 0.5).

## Results

### Spatial-only model

In the spatial-only model, species’ traits were strong predictors of survival probability and were consistent in direction across global, regional and local spatial scales (Fig. 3, Table S1, Table S2, Table S3). Extinction risk was higher for species with large body masses, small brains, carnivorous diets, long generation lengths, or terrestrial (rather than scansorial) habits. Scansoriality was non-significant at the local scale, though its direction – scansorial species being less vulnerable than terrestrial species– was consistent with those at the global and regional scales.

**Figure 3.**
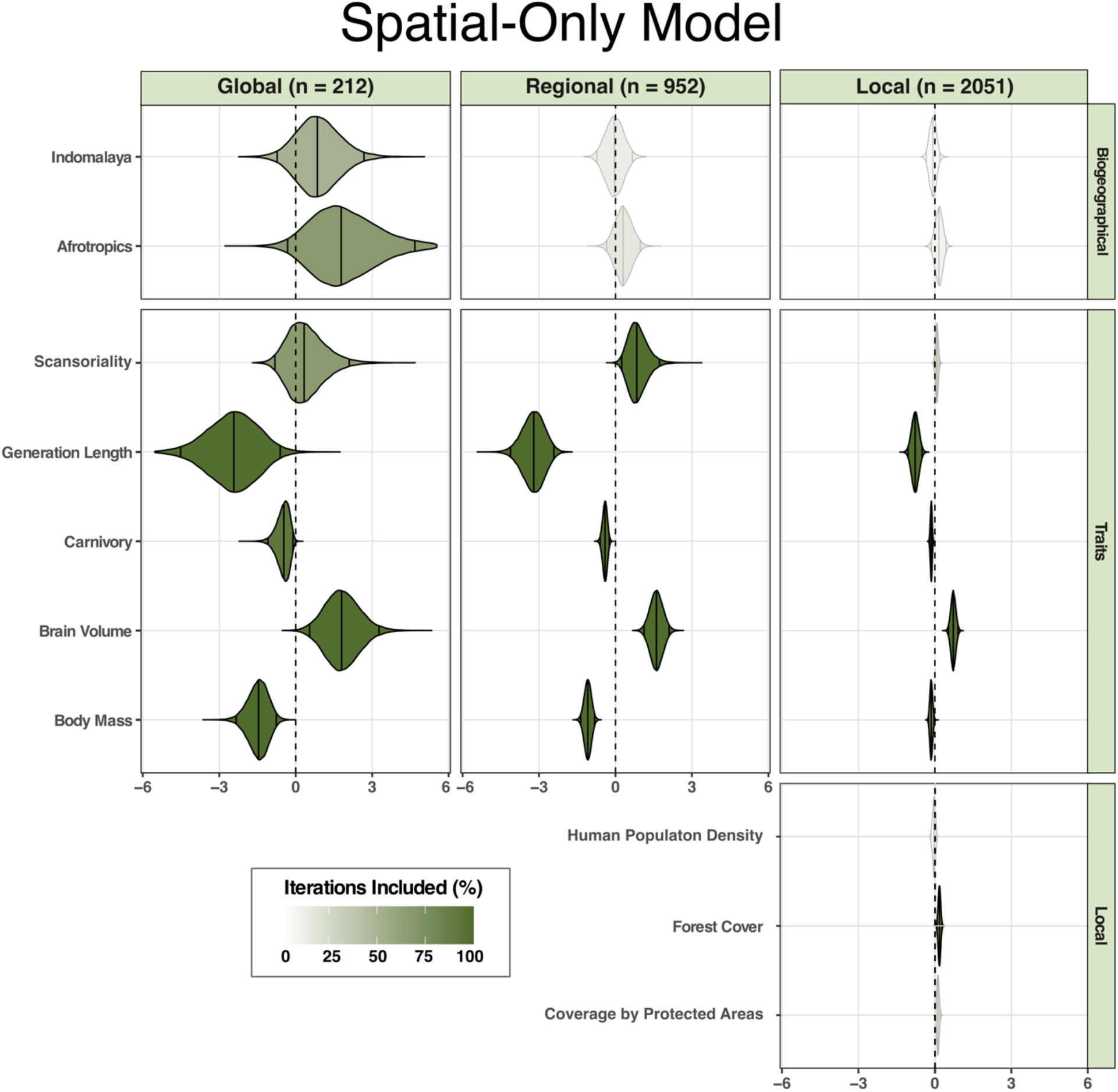
Predictors of extinction risk were similar across the global, regional and local scales when regressions only varied in their spatial resolution. Violin plots show posterior distributions of coefficient estimates of the spatial-only model. Vertical lines inside the violin plots show the 95% credible intervals and median values. The color intensity indicates how often each predictor was included in the model. Variables included in more than 50% of the iterations are drawn with black borders. Variables included in fewer than 50% of the iterations are drawn using gray borders. The Neotropics are the intercept level to which the other biogeographical realms are compared.

Species survival probabilities also varied depending on where species occurred. First, species from different biogeographical realms had different survival probabilities on the global scale; Afrotropical and Indomalayan species had higher survival probabilities than Neotropical species, with the highest survival in the Afrotropics. Second, at the local scale, species were more likely to survive in areas with a high forest cover, whereas human population density or protection status of sites did not significantly influence survival probabilities.

### Spatial-temporal model

In the spatio-temporal model, predictors of extinction risk were not consistent across scales (Fig. 4, Table S4, Table S5, Table S6). At each scale, predictors differed in direction and magnitude. At the global, long-term scale, results resembled the spatial-only model; extinction was more likely for species that were large, had long generation lengths, small brains, or carnivorous diets. At the regional scale, small species and scansorial species were more likely to survive, but no other traits were significant. However, at the local, short-term scale, large species and species with long generation lengths were more likely to survive, opposite to findings at the global, long-term scale.

**Figure 4.**
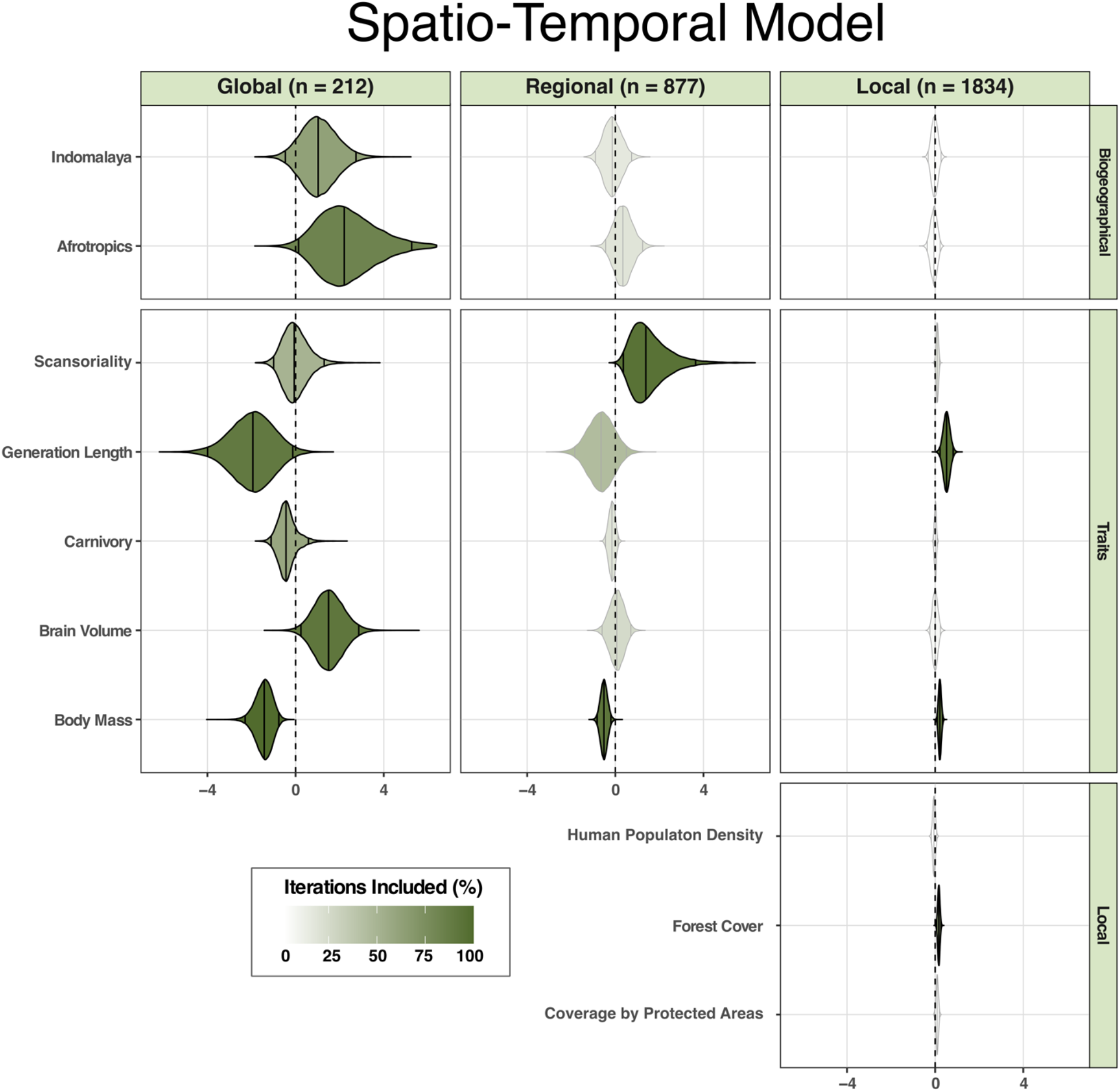
Predictors of extinction risk varied across the global, regional and local scales when regressions vary in their spatial resolution and temporal extent. Violin plots show posterior distributions of coefficient estimates of the spatio-temporal model. Vertical lines inside the violin plots show the 95% credible intervals and median values. The color intensity indicates how often each predictor was included in the model. Variables included in more than 50% of the iterations are drawn with black borders. Variables included in fewer than 50% of the iterations are drawn using gray borders. The Neotropics are the intercept level to which the other biogeographical realms are compared.

Biogeographical differences in survival probability were similar to those in the spatial-only model. At the global scale, Afrotropical and Indomalayan species had higher survival probabilities than Neotropical species, and the effect was strongest in the Afrotropics. Likewise, at the local scale, species were more likely to survive in areas with higher forest cover.

### Phylogenetic models

We also ran both models with a correction accounting for the phylogenetic relationships between species (see Methods and SI Models with Phylogenetic Correction). In both cases, phylogeny was a significant predictor of survival probability at all three scales (Fig. S1, Fig. S2). However, adding a phylogenetic correction to the model did not meaningfully change the effect of other predictors, except for carnivory which was strongly correlated with phylogeny (Fig. S5).

## Discussion

For large-scale studies to inform conservation, their findings must be relevant to the often narrower spatial and temporal scales at which practitioners and policymakers work [4]. We studied the degree to which the predictors of extinction risk were consistent across scales for tropical forest mammals. We found that intrinsic predictors of extinction risk were similar across the global, regional and local scale when regressions only varied in their spatial resolution and used the same 130,000-year temporal extent (spatial-only model). Under this model, mammal species with longer generations, carnivorous diets, smaller brains, or larger bodies were more likely to go extinct at all spatial scales. In contrast, extinction risk predictors were variable when the temporal extents of the regressions became increasingly shorter (spatio-temporal model). While the spatio-temporal model identified similar predictors as the spatial-only model at the global scale, it found little consistency between large versus small spatial scales, even finding opposite effects in the case of body mass and generation length. This discrepancy between scales and models suggests that the processes driving extinction at the local scale today may differ from those that have been driving large mammal extinctions over the past 130,000 years.

Moreover, these contrasting results highlight how different temporal extents provide different perspectives on these changing drivers of extinction. Large-scale extinction risk studies have been criticised for their inconsistent findings and their lack of clear conservation messages [16]. Our results suggest that this variation amongst study findings may partly arise due to differences in their temporal extent, which may lead to opposite conclusions. For example, studies of brain size and its effect on extinction risk that only include extant species have found that extinction risk is higher for species with large brains [30,31]. In contrast, studies that also include globally extinct species, including taxa from the Pleistocene or earlier periods, found that extinction risk is lower for species with large brains [21,32]. Nonetheless, the temporal extent of studies often remains implicit, and has been less examined than the spatial and taxonomic extent (for example, see Cardillo and Meijaard [16] who discuss the influence of spatial and taxonomic but not temporal extents on studies of extinction risk).

Temporal extents change the observable ecological patterns in studies of extinction because they affect the species pool (and thereby also taxonomic extent). By only considering recent temporal extents, which can happen due to shifting baselines [17], studies of extinction exclude species that went extinct previously. However, because extinctions are non-random, such species are more likely to have extreme trait values, such as large body sizes or unique diets [9,33]. The surviving species pool will, therefore, possess properties that differ from those of the original species pool, and inferences based on one may not apply to the other [34]. Specifically, studies with short temporal extents only view a limited range of the possible set of trait values, which can dampen the observable effect of processes that caused extinction in the past. Short temporal extents can, therefore, hide long-term drivers of extinction or underestimate their importance.

The reverse may also be true. Including long-extinct species can overshadow contemporary trends affecting the remaining species, either because vulnerable species have been removed or because the drivers of extinction have changed. For example, our spatio-temporal model predicted that long-lived, large-bodied species were more likely to survive at the local, short-term scale. We hypothesize this pattern exists because camera trap data (including those in our study) predominantly come from protected areas [12,35]; these are sites where threatened species receive special protection and where large mammal population trends tend to be more positive [35]. We did not detect similar effects in the spatial-only model. The strong negative effects that body size and generation length have historically had on extinction likely overrode the more subtle contemporary correlation between large mammals and survival in protected areas.

Conservationists can, thus, use macroecological findings to identify threatened species if they are aware of the extent of the studies they rely on (temporal or otherwise). However, uncritically adopting macroecological findings can cause one to misidentify the determinants of extinction risk at smaller scales. Long-term extents will give greater emphasis to trait values that have already been lost, which can be valuable when highlighting the functional gaps in assemblages and guiding functional restoration efforts. Yet, emphasising extinction predictors that are no longer relevant may prevent conservationists from accurately identifying present-day threatened species. Studies with short temporal extents are, arguably, better at identifying present-day risks. However, if they exclude long-extinct species, some predictors of extinction may also be missed. For example, imagine an extinction driver is ongoing but most species vulnerable to the driver have perished and are forgotten. Yet, a few species vulnerable to the driver survived in the community (e.g. due to random chance). Such species risk being classified as “not vulnerable” because there is no recent evidence that their traits predispose them to extinction. Analysing extinction risk from both short and long-term perspectives may, ultimately, provide the most comprehensive view of extinction risk, as it can highlight changes in the relative importance of different drivers of extinction.

### Do our findings apply to other taxa, ecosystems or predictors?

The findings of our spatial-only model concur with well-known macroecological patterns of extinction in vertebrates, such as the fact that species with large body sizes, long generation lengths and comparatively small brains have a higher extinction risk [13,21,36,37]. Likewise, we identified the Afrotropics as having the fewest global extinctions, followed by Indomalaya and finally the Neotropics, likely reflecting species’ adaptation or co-evolution with humans in areas that have a long history of human presence [38]. As reported in previous studies, our results also suggest that carnivores are comparatively more threatened than other dietary groups ([39]; but see [40]). Because these patterns are consistent with studies on other taxa and habitats, we suspect our findings will apply to vertebrates more broadly, and not just terrestrial forest mammals.

Scale generalisability may not be equal for all processes, however. We studied a process with clear links between scales, where global extinctions result directly from extinctions at smaller scales. Scale generalisability may be lower in other processes. For example, research suggests species’ habitat selection may be difficult to generalise, because the act of selecting habitats happens on one or only a few scales. Certain scales may, therefore, be more informative than others [41].

Additionally, scale generalisability may be lower for extrinsic predictors of extinction risk. Our study primarily compared similarities between species’ trait variables, but extrinsic variables, such as forest cover, economic status, or cultural attitudes are equally important for conservation issues. It is, arguably, unlikely that all extrinsic predictors would exhibit similar patterns across scales. In fact, the one extrinsic variable we compared across scales (biogeography) did not display scale generalisability.

Finally, our data predominantly come from protected areas, reflecting available camera trap data [12]. This could influence the extinction predictors we identify. Non-protected areas often experience pressures that are more intense or unlike those found in protected areas, such as agricultural expansion, urbanisation, hunting or resource extraction [42]. Consequently, it is unclear whether the positive trends in the spatio-temporal model’s local regression are the result of scale differences or of the data at our disposal. This uncertainty does not change our conclusion, however, because the contrasting findings of both models still illustrate how changes in temporal extent can change a study’s findings. Nevertheless, the higher survival of species with traits that are normally associated with extinction risk does raise questions about the influence of protected areas on biodiversity data. Possibly, such findings indicate that protected areas can safeguard vulnerable species, as was suggested for species with large body sizes [35]. Yet, such patterns could also arise due to selection biases in protected area placement or extinction debts [42].

More studies are, therefore, needed, that analyse to what degree studies with large spatial or temporal extents can inform conservation. Ideally, such studies should analyse different processes, different types of predictors, different study sites (including non-protected areas), and taxonomically diverse groups. We specifically call for more research regarding the effect of large temporal extents on local-scale studies. Many conservation actions take place at the local scale, but we found that temporal extents could fundamentally change the conclusions of local-scale studies on which such actions often rely. Ultimately, we hope that such studies will help clarify which macroecological findings are relevant for more local conservation efforts and guide more effective conservation strategies.

## Methods

### Site Selection

We assembled a comprehensive list of camera trap locations (n = 337) across the pantropics, situated inside wet tropical forests of varying protection status. We collated data from the TEAM camera trapping protocol [43], Amazonia CAMTRAP [44], as well as individual datasets provided by authors of this manuscript (e.g. [45]). First, we manually eliminated datasets outside forest areas (n = 3). We also removed a single site on Madagascar. Madagascar is biogeographically distinct and we did not want to introduce sampling bias by representing this region with a single site.

To eliminate duplicate records and minimise spatial autocorrelation among sites, we calculated 10 km buffers around each camera trap. Camera traps with overlapping buffers were then clustered together, resulting in 140 camera trap clusters (henceforth, sites) that were at least 20 km apart. We visually inspected these clusters to ensure they were clearly separated.

The detection of species is not constant between cameras and between sites [46]. Therefore, we assessed the species accumulation curve at each site, retaining only sites where the curve converged to a slope of 0.2 or lower (indicating that most species present at the site and detectable by camera traps had been detected; see SI SI_Data_1_rarefaction_curves.pdf). This process yielded a final selection of 64 sites, 46 in the Neotropical realm, 10 in the Afrotropical realm, and 8 in the Indomalayan realm. Due to the clustering methodology, our study sites could be entirely within, partially inside or outside protected areas (as defined by the World Database on Protected Areas). However, sites that are either partially or fully within protected areas are more common in our dataset (n = 56, Fig. S3), as camera traps are still primarily deployed in protected areas and other areas of conservation interest [12,35].

### Species Lists

For each site, we created four species lists: 1) present-natural, 2) surviving globally, 3) surviving regionally, and 4) surviving locally (Fig. S4). These species lists are nested: If a species occurs in a given species list it also occurs in the species lists above it. We used the different species lists to create the presence-absences used in the model regressions (see Statistical Model).

The **present-natural** species list is the total species pool. It is based on the species list of PHYLACINE v1.2.1 [20] rather than camera-trap data, and contains all species expected to occur at a site if there had been no local or global anthropogenic extinctions since the Eemian period, which started ±130,000 years ago. The Eemian period provides a reference for a world with a comparable climate and species composition but with limited human impacts, and is commonly used in macroecological studies of extinction (e.g. [18]). We created the present-natural species lists by overlaying the present-natural ranges from PHYLACINE v1.2.1 [20] with the locations of each site.

Present-natural ranges are not past ranges. They estimate where species would occur today in the absence of human impacts. For most species (n = 156), these ranges are identical to their IUCN ranges. For species with human-caused range modifications (n = 54, including global extinctions), the present-natural ranges predict the distribution without anthropogenic modifications [20]. For globally extinct species, the present-day range is estimated using a variety of methods, but in most cases, is based on the present-day occurrence of species with which the extinct species co-existed (for details, see [20]).

To keep the present-natural species list comparable to the surviving locally species lists which is based on camera trap data (see below), we excluded species with primarily aquatic or arboreal lifestyles (though we retained scansorial species, i.e. largely terrestrial species with the ability to climb), as well as species smaller than 1 kg, because they are difficult to detect with ground-based camera traps [47]. We also excluded extinct human species because their interactions with *Homo sapiens* were likely different from those of non-human mammals.

Due to the coarse resolution of present-natural maps (96.5 × 96.5 km at 30° North and South), our species list included species that would not have occurred in the tropical rainforests of our study sites. Therefore, we checked the ecology of each remaining species and excluded species that would not occur in tropical rainforests (n = 260, see SI SI_Data_2_excluded_species.csv). We relied on the handbook of the mammals of the world series where possible [48–51], and otherwise on other scientific and gray literature. We paid particular attention to globally extinct species, and specifically excluded grazers and species tied to open ecosystems.

There were 57 (out of 2051) cases where camera traps recorded extant species at a site even though they were not predicted to occur there by PHYLACINE’s present-natural maps. We added these species to all relevant species lists and treated them as if they had been predicted to occur there. Most of these inconsistencies were the result of taxonomic differences between datasets or IUCN ranges that underestimated species’ ranges and were subsequently copied by PHYLACINE (SI SI_Data_3_range_inconsistencies.csv).

Due to uncertainties in the (sub)fossil record, it is possible that some undocumented extinctions have occurred over the last 130,000 years, especially in tropical forests which do not preserve fossils well [52]. Some globally extinct species may, therefore, still be missing from our present-natural species pool. Nonetheless, the extinctions of Quaternary medium-large mammals are well-known compared to most other species groups and periods, due to its recency and the fact that mammals are amongst the most-studied groups in palaeo-archaeological sciences [53].

The **surviving globally** species list for a given site contains all species from the sites’ present-natural species list that remain globally extant. Out of the 210 species, 199 remain globally extant and 11 have gone globally extinct.

The **surviving regionally** species list for a given site contains all species from the sites’ surviving globally species list that are still extant in the region in which the site is located (Fig. 1). We created the surviving regionally species list by overlaying IUCN ranges (Version 2022-1, using the extant, probably extant, possibly extant parts of the range) with the “Basic Recording Units’’ from the World Geographical Scheme for Recording Plant Distributions (WGSRPD) [54]. “Basic Recording Units” (which we call “regions”) are approximately evenly sized geographical entities delineated by political borders and coastlines. They represent areas that broadly share a biogeographical history, and fall under the same administrative entities, meaning they are subject to similar conservation policies. In the Virunga Massive, the camera trap set-up straddled the border between Rwanda and Uganda, thus overlapping with two Basic Recording Units. However, we assigned Rwanda as the basic recording unit as 55 out of 60 camera traps were located in Rwanda and as all species recorded on the Ugandan site were also recorded on the Rwandan side. Species were considered surviving in the state if their IUCN range maps overlapped with the “Basic Recording Units” in which the site was located.

The **surviving locally** species list contains all species extant at a site. We listed species as locally surviving if they had been observed at the site according to our camera trap records. We removed all records of humans, domestic species, and species smaller than 1 kg. Moreover, we removed records of primarily aquatic or arboreal species (though we retained scansorial species). Removing these species left us with 142 species that were observed at one or more of our sites.

Note that our method assumes that a species should occur at a site if that site falls within the species’ present-natural range. However, species’ ranges are seldom completely occupied, and while likely, we cannot be certain that a given “predicted but absent” species would have occurred at any given site. The local scale “extinctions” in our study are therefore, “presumed extinctions”, which potentially inflates the number of “extinct species” at the local scale.

### Predictor Variables

We collected several potential predictors of extinction and divided them into the following groups: *biogeographical variables, species traits, site variables*, and *phylogenetic variables. Biogeographical variables* describe biogeographical differences between regions. *Species Traits* describe properties of species (e.g. body mass). *Site variables* describe properties of the site (e.g. forest cover). *Phylogenetic variables* describe the phylogenetic relationships between species.

#### Biogeographical Variables

To account for biogeographical differences in extinction risk, we assigned each site to one of three realms with unique biogeographical histories: the Afrotropics, Indomalaya, and the Neotropics. We included the biogeographical realm as a potential predictor of extinction at each scale. Biogeographical realm operates like a “blocking factor”. It accounts for differences in extinction risk between biogeographical areas that we cannot explain with our other variables.

#### Species Traits

For all 210 species in our study, we collected trait information on: body mass (kg), vertebrate carnivory (%), scansoriality (terrestrial | scansorial), generation length (days), and brain endocast volume (ml). All these traits have been suggested as predictors of extinction by other studies [21,36,55,56]. Body mass is a proxy for body size, one of the most fundamental correlates of a species life-history [57]. Vertebrate carnivory reflects a species diet and trophic level. Here, it is defined as the percentage of vertebrate consumption in the diet, as recorded by PHYLACINE v1.2.1 [20]. Due to how it is defined, this variable is almost entirely the inverse of a species’ plant consumption, and thus allows us to differentiate between herbivores, omnivores, and carnivores. Scansoriality reflects a species’ arboreality, and thus its ability to avoid ground-based human hunters, and to some extent its dependence on tree cover [36]. Generation length reflects species demographic rates, and correlates with how quickly species can recover from low population sizes. Brain volume affects extinction risk, as large (relative) brain volumes give species the behavioral flexibility to deal with threats and novel environments [21,32,58], but it also requires large energetic allocations that could increase extinction risk [30,31].

We collected body mass and diet estimates from PHYLACINE [20]. We collected scansoriality from Lundgren et al. [24] and Wilman et al. [25] and completed data gaps by manually reviewing the ecology of species with missing arboreality data. We collected endocast volumes from Dembitzer et al. [21] or derived them from brain mass estimates in Burger et al. [22]. We converted brain mass to endocast volume, following the method in [21]. First, we converted brain mass to brain volume by dividing it by 1.036. Next, we converted brain volume to endocast volume using the formula: *log(Endocast Volume) = − 0*.*0015 + 1*.*0222 * log(Brain Volume)*. We collected generation length from Pacifici et al. [23]. We phylogenetically imputed missing traits (Table S7) using the PHYLACINE v1.2.1 phylogenies, and the Rphylopars package [59] in R version 4.2.2 (2022-10-31). We imputed all missing traits of all non-flying, terrestrial mammals (n = 4535 species) using 1000 different phylogenetic trees, and using mass, diet, scansoriality, endocast volume and generation length as predictors. We, thus, had 1000 imputed estimates for each missing trait value, which were later provided to the model (see Statistical Model).

To emphasise the relative size differences between species we used the logarithm of body mass rather than absolute body mass values in our analysis. To avoid multicollinearity between body mass, brain volume and generation length, we regressed log-transformed body mass against log-transformed brain volume and against log-transformed generation length using a simple linear model. We used the residuals of these regressions as input variables in our statistical model. These residuals model the effects of brain volume and generation length on extinction risk that cannot be explained by body mass. The subsequent collinearity between variables is reported in Fig S5 and Fig. S6.

Species’ traits can vary across space and time. However, the traits used in this study are robust and unlikely to be strongly affected by such variability. For example, our diet estimates broadly differentiate between carnivores, omnivores and herbivores. Such classifications would not change over the 130,000 years considered in this study, or across different sites. Similarly, most modern mammal species have existed for several 100,000s to millions of years [60], indicating that major morphological traits (e.g. body size or scansoriality) have remained consistent over timescales that exceed those examined in this study.

#### Site Variables

Our model included three site variables: degree of forest cover, human population density, and the percentage of camera traps at a site located inside protected areas. We estimated forest cover by using the Global Forest Change (GFC) dataset [26], which contains canopy cover data for the year 2000 (30 × 30 m resolution), as well as data on net changes in canopy cover between 2000 and 2012. First, we generated a forest cover map for the year 2000 (defining forest as tree canopy cover >75%, and masking water bodies). Next, we calculated the net change from 2000-2012 and used it to create forest cover maps for the year 2012. We then calculated the forest cover in 2012 in a 5, 10, and 20 km buffer around each site. We extracted population density from CIESIN’s Gridded Population of the World (v4.11) dataset [27]. This dataset maps human population density across the planet (30 arc-second resolution) in 5-year timesteps from 2000 until 2020. We calculated the median human population density in 2010 in a 5, 10, and 20 km buffer around each site. We log-transformed the data before putting it in the model. The main models used the 10 km buffer, but to ensure that model findings were stable, we also ran models using the 5 and 20 km buffers. These did not meaningfully change the results (Fig. S7, Fig. S8, Fig. S9, Fig. S10).

#### Phylogenetic Variables

We ran alternative versions of each model (SI *Models with Phylogenetic Correction*), using phylogenetic eigenvectors to account for the phylogenetic relationships amongst species [61]. We used Rphylopars [59] to calculate the phylogenetic distances between all species based on 1000 phylogenetic trees from PHYLACINE [20]. We performed a principal coordinate analysis (PCoA) on each of the resulting distance matrices. We averaged the 1000 PCoA outputs and included the first two axes of the averaged output in our model as explanatory variables, which explained 39.19% of variation.

### Statistical Model

We developed a Bayesian model to assess the survival probability of all 210 mammal species (see supplement Vignette.html) across all 64 sites (Fig. 1) and across three different scales (global, regional, local). The Bayesian models consisted of three logistic regressions, corresponding to each scale (Fig. 2). The global regression modeled the variables that predict whether species survive globally. The regional regression modeled the variables that predict whether species survive regionally. The local regression modeled the variables that predict whether species survive locally.

*The* survival probability of species *sp* at scale *sc* can be expressed as the outcome of a regression:

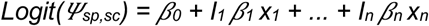

*where β*_*0*_ *is the intercept, β*_*1-n*_ are the coefficients of predictor *i, and x*_*i*_ is the value of predictor *i*. Indicator variables *I*_*i*_ denote inclusion (*I*_*i*_*=1*) or omission (*I*_*i*_*=0*) of a given coefficient and associated predictor in the regression. The inclusion variables are sampled from a Bernoulli distribution with inclusion probability parameter *p*_*i*_

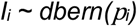

Parameter inclusion probability estimation is implemented as part of reversible jump MCMC sampling *(rj MCMC) to allow sampling of the entire parameter space and proper model convergence*.

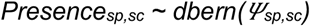

The presence of species *sp* at scale *sc (Presence*_*sp,sc*_) is then estimated using a Bernoulli distribution.

We ran two different versions of the model (Fig. 2). In the spatial-only model, the only difference between regressions is their spatial resolution (Table 1). In the spatio-temporal model, the spatial resolution changes as well as the temporal extent (Table 2). We do so by excluding species from smaller scales if they have already gone extinct at a higher scale, broadly corresponding to the chronology of mammal extinctions and the subsequent absence of species from scientific studies due to shifting baselines [17]. In the spatio-temporal model, the global, regional and local scales, therefore, correspond to studies with long, intermediate and short temporal extents, respectively. Excluding these species gives each regression different but nested species pools. Note that both models are identical at the global scale and that all regressions have the same pantropical extent.

**Table 1.** The Spatial-Only Model and the species lists they use in each regression.

|  | **Total Species Pool** | **Present Species** |
| --- | --- | --- |
| **Global** | Present-Natural | Surviving Globally |
| **Regional** | Present-Natural | Surviving Regionally |
| **Local** | Present-Natural | Surviving Locally |

**Table 2.**
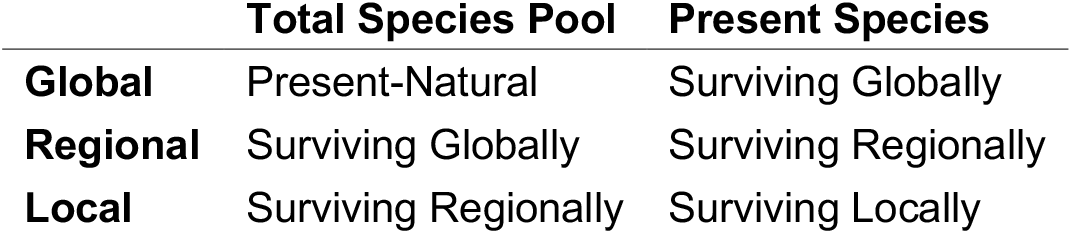
The Spatio-Temporal Model and the species lists they use in each regression.

|  | **Total Species Pool** | **Present Species** |
| --- | --- | --- |
| **Global** | Present-Natural | Surviving Globally |
| **Regional** | Surviving Globally | Surviving Regionally |
| **Local** | Surviving Regionally | Surviving Locally |

We fitted the model using the Nimble package v0.12.2 [62] in R, and ran 6 chains over 50000 iterations, with a burn-in period of 1000 iterations and a thinning interval of 5. We used the reversible-jump Markov Chain Monte Carlo (MCMC) method to highlight the most significant predictors in each regression [29]. The reversible jump method associates each model coefficient with a binary inclusion parameter (0: excluded, 1: included). The model estimates the likely value of this model parameter over all model iterations and provides an estimate of the inclusion probability. The inclusion probability is the proportion of times a variable is included in the model across all MCMC iterations. It is a metric of the evidential support for a given predictor. We considered variables significant if they had an inclusion probability of 0.5 or higher, meaning they were more likely to be included rather than excluded by the model.

We accounted for the uncertainty surrounding missing trait values by, for each missing trait value, providing all 1000 imputed trait values as a data input to the model. Based on these 1000 imputed values, the model estimated the underlying distribution for each missing trait value and each MCMC iteration and randomly drew values from that distribution. This meant that our total model estimated its parameters using different potential trait values, but that all three regressions used identical trait values during each MCMC iteration, keeping their results comparable at each step.

All variables were included at all scales, except for site variables. Site variables were included at the local scale only to account for their known effects on extinction risk rather than to test their consistency across scales. For an overview of all variables included in each regression, see Table S1, Table S2, Table S3, Table S4, Table S5, and Table S6. We scaled all predictor variables prior to the analysis, by centering the data around its mean and dividing it by its standard deviation. We used uniform distribution ranging from 0-1 as the priors for the reversible jump parameters. We used a logistic distribution with location 0 and scale 1 as the prior for the coefficient parameters.

We calculated the median of the posterior distribution of each coefficient, the 95% Bayesian credible interval, and inclusion probability. We made sure the model parameter estimates converged, by visually appraising the convergence plots and by checking the Gelman-Rubin statistic for each parameter ensuring they remained below 1.1.

## Supporting information

Vignette.html

SI_Data_1_rarefaction_curves.pdf

SI_Data_2_excluded_species.csv

SI_Data_3_range_inconsistencies.csv

SI_Data_4_Trait_Array.rds

SI_Data_5_Species_Traits_and_Metadata.csv

SI_Data_6_site_info.csv

SI_Data_7_species_park_list.csv

## Acknowledgments

We thank Rasmus Østergaard Pedersen for his help regarding phylogenetic imputations, and Rhys Taylor Lemoine for his help regarding biogeography. We thank Soumen Dey for his help with Nimble and Bayesian analyses. We thank Arafat Mtui, Julia Salvador, and the many people who helped to collect the camera trap data used in this study. We also thank Zimices, Sarah Werning, Steven Traver whose silhouette artwork we used (under CC BY-NC 3.0), and Charles J. Sharp whose picture we used to create the silhouette of Rusa unicolor (under CC BY-SA 4.0).

