## Supplementary Information for "Predictors of extinction risk in large tropical forest mammals: from global to local"

Clarification of the Trait Metadata

Here, we summarise the different terms that can be found in the trait metadata (see SI SI_Data_2_excluded_species.csv). To the extent possible, we followed the metadata classifications used in the primary, original sources. For a detailed description of the metadata we refer to the original sources.

**Mass.Method**

*Metadata taken from Faurby et al. (2018)* [25]

- **Reported:** Reported by other source
- **Assumed isometric based on X:** Isometrically estimated based on a certain known morphological trait (e.g. mandible size, skull size, bone size, shoulder height, head-body length, etc.)
- **As relative of suggested similar size:** The size is known to be similar to that of a related species.
- **Phylogenetically Imputed:** phylogenetically imputed by this study

**Diet.Method**

*Metadata taken from Faurby et al. (2018)* [25]

- **Reported Observed Species:** The data was taken directly from the source, providing numerical data. The data is based on direct observations. The data is reported at the species level.
- **Reported 000 Species:** The data was taken directly from the source, providing numerical data. The source does not specify which method was used to collect the data. The data is reported at the species level.
- **Estimated Observed Species:** The data been manually estimated by the authors of PHYLACINE, based on the listed evidence. The data is based on direct observations. The data is reported at the species level.
- **Estimated Expert Species:** The data been manually estimated by the authors of PHYLACINE, based on the listed evidence. The data is based on classifications by experts. The data is reported at the species level.
- **Estimated Expert Genus:** The data been manually estimated by the authors of PHYLACINE, based on the listed evidence. The data is based on classifications by experts. The data is reported at the genus level.
- **Estimated Craniodental Species:** The data been manually estimated by the authors of PHYLACINE, based on the listed evidence. The data is based on craniodental measurements. The data is reported at the species level.

- **Estimated Craniodental Genus:** The data been manually estimated by the authors of PHYLACINE, based on the listed evidence. The data is based on craniodental measurements. The data is reported at the genus level.
- **Transformed 000 Species:** The data was taken directly from the source, providing numerical data, but have been mathematically transformed to conform to the PHYLACINE format. The source does not specify which method was used to collect the data. The data is reported at the species level.
- **Transformed 000 Genus:** The data was taken directly from the source, providing numerical data, but have been mathematically transformed to conform to the PHYLACINE format. The source does not specify which method was used to collect the data. The data is reported at the genus level.
- **Phylogenetically Imputed:** phylogenetically imputed by this study, at the species level.

**Generation.Length.Method**

*Metadata taken from Pacifici et al. (2013)* [46].

- **GMA:** value taken from IUCN Red List data.
- **Rspan-AFB:** Generation length calculated as the difference between reproductive life span and age at first birth.
- **Rspan-AFR(SM+Gest):** when data on age at first reproduction were not available, the source calculated this parameter as the sum between age at female sexual maturity and gestation length.
- **Rspan-ASMmales:** Generation length calculated with age at sexual maturity for males, when data on age at first reproduction for females were not available.
- **Phylogenetically Imputed:** phylogenetically imputed by this study.

**Endocast.Method**

*Metadata taken from Dembitzer et al. (2022)* [34].

- **Volume:** Endocast volume taken directly from the reported source.
- **Mass:** Endocast volume was converted from Brain mass following the method used in Dembitzer et al. (2022).
- **Phylogenetically Imputed:** phylogenetically imputed by this study.
- **Method not provided by original source:** The original source did not specify how the values were derived.

**Stratum.Method**

*Metadata taken from Lundgren et al. (2021)* [43] *and Wilman et al. (2014)* [44].

- **Reported:** Stratum use reported by another source.

- **Functional morphology**: Stratum use inferred from the functional morphology of the species.
- **Expert Judgement:** Informed judgement based on the known ecology of the species.

**Observed X:** The species has been observed to use a given stratum.

<u>Models with Phylogenetic Correction</u>

In both the spatial-only and the spatio-temporal model, phylogenetic relationships were significant at all three scales (Fig. S1 and Fig. S2). Species with high values on phylogenetic dimension 1 (Fig. S11) were mostly marsupials or rodents (e.g. Didelphis, Dasyprocta). Species with low values were mostly carnivorans (e.g. Leopardus, Prionailurus, Puma). Species with high values along dimension 2 tended to be ungulates (e.g. Cephalophus, Mazama).

Including these phylogenetic variables in the model had no strong influence on other variables, except for carnivory. The strong correlation between carnivory and phylogenetic dimension 1 led to some convergence issues when both variables were included in the model, suggesting that both variables capture similar information. The issues disappeared when either one of the variables was excluded.

We note that phylogenetic relationship between species are used for two different purposes in our study: imputing trait values and the phylogenetic correction. Theoretically, this could inflate the phylogenetic relationships we find and weaken the effect of non-phylogenetic predictors. However, this is not what we find. While we do find a phylogenetic effect on extinction risk, it is not extremely high, nor does it greatly influence other predictors.

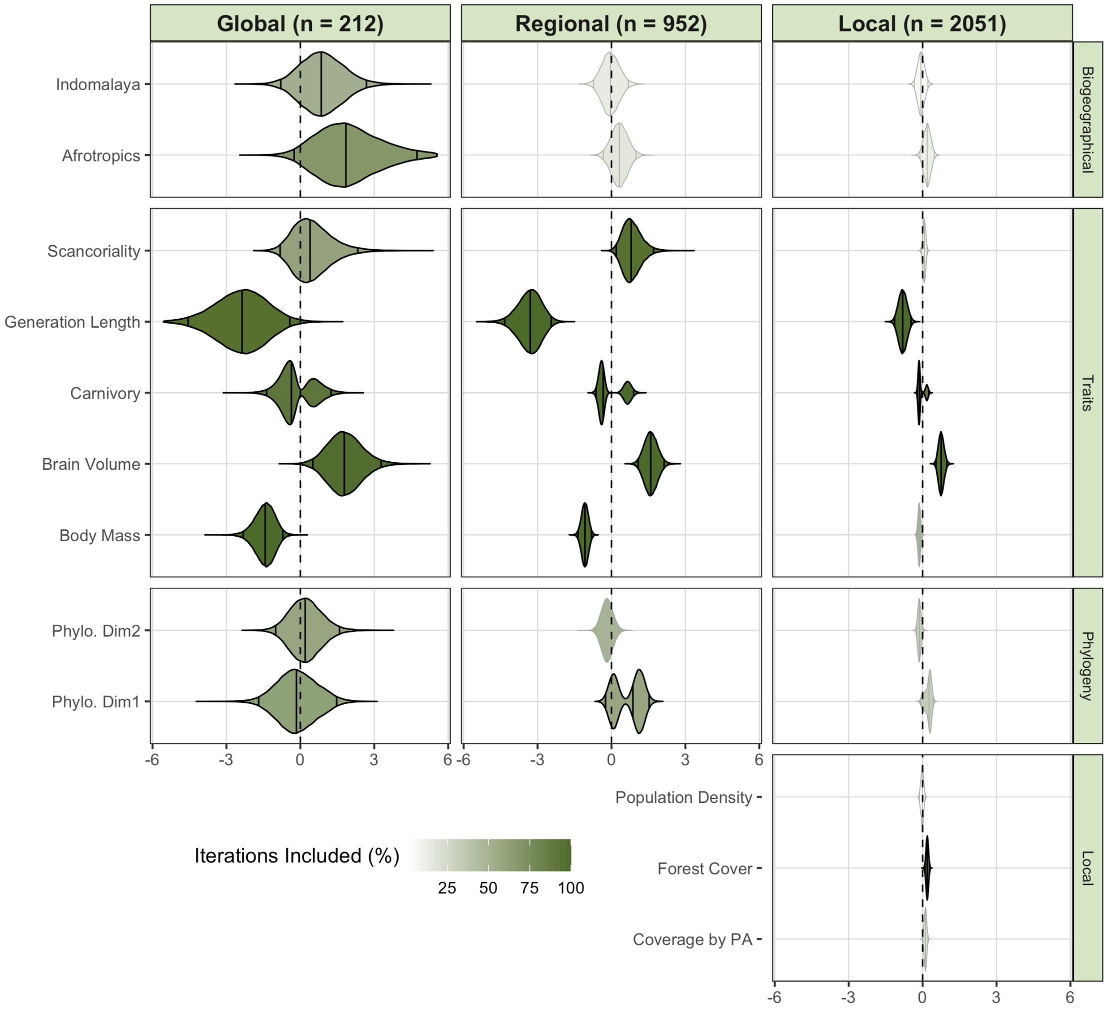


**Figure S1. Posterior distributions of the coefficient estimates of the spatial-only model with phylogenetic correction.** Vertical lines inside the violin plots show the 95% credible intervals and the median values. The colour intensity of the plots show how often each variable was included in the model. Variables included in more than 50% of the iterations were drawn with black borders. Variables included in fewer than 50% of the iterations have been drawn using grey borders. The Neotropics are the intercept level to which the other realms are compared.

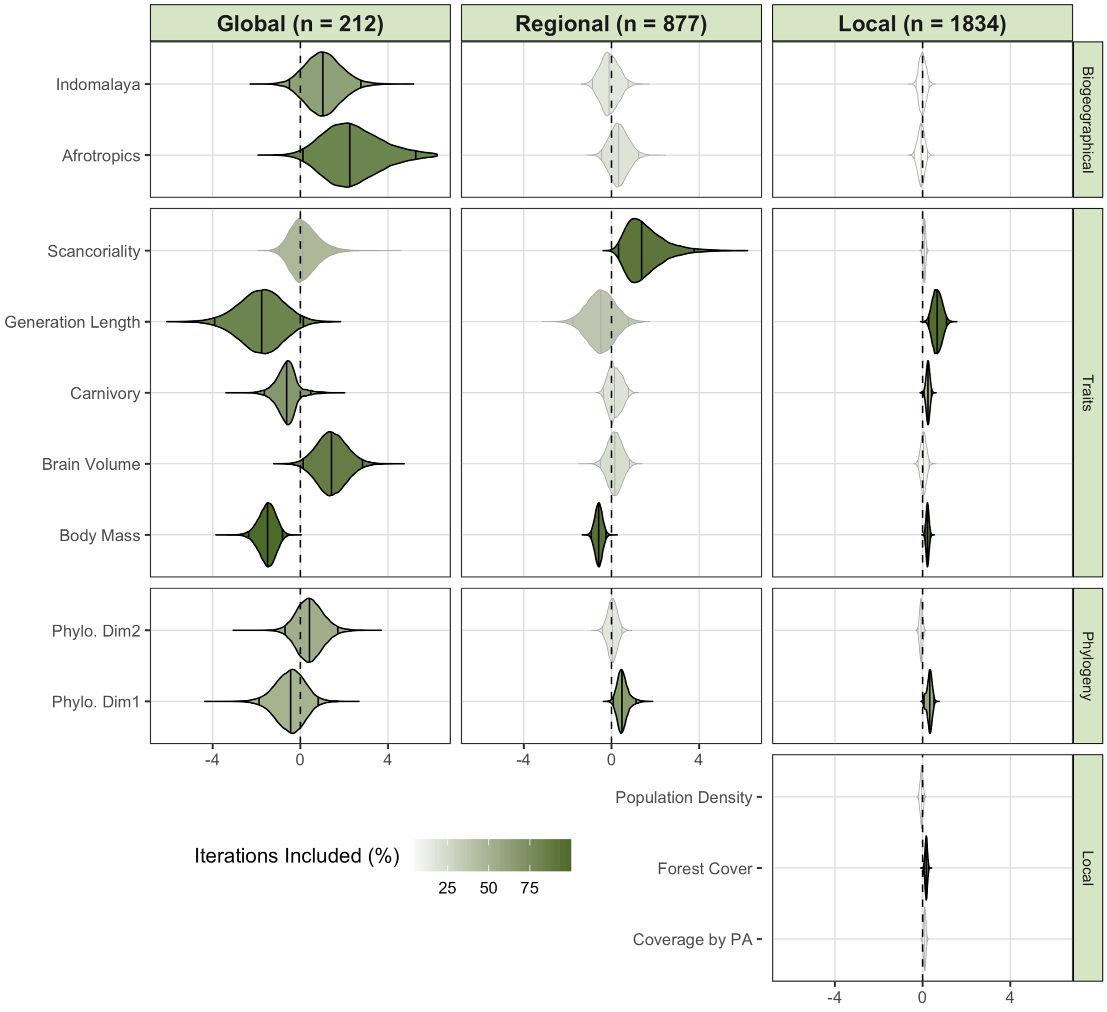


**Figure S2. Posterior distributions of the coefficient estimates of the spatio-temporal model with phylogenetic correction.** Vertical lines inside the violin plots show the 95% credible intervals and the median values. The colour intensity of the plots show how often each variable was included in the model. Variables included in more than 50% of the iterations were drawn with black borders. Variables included in fewer than 50% of the iterations have been drawn using gray borders. The Neotropics are the intercept level to which the other realms are compared.

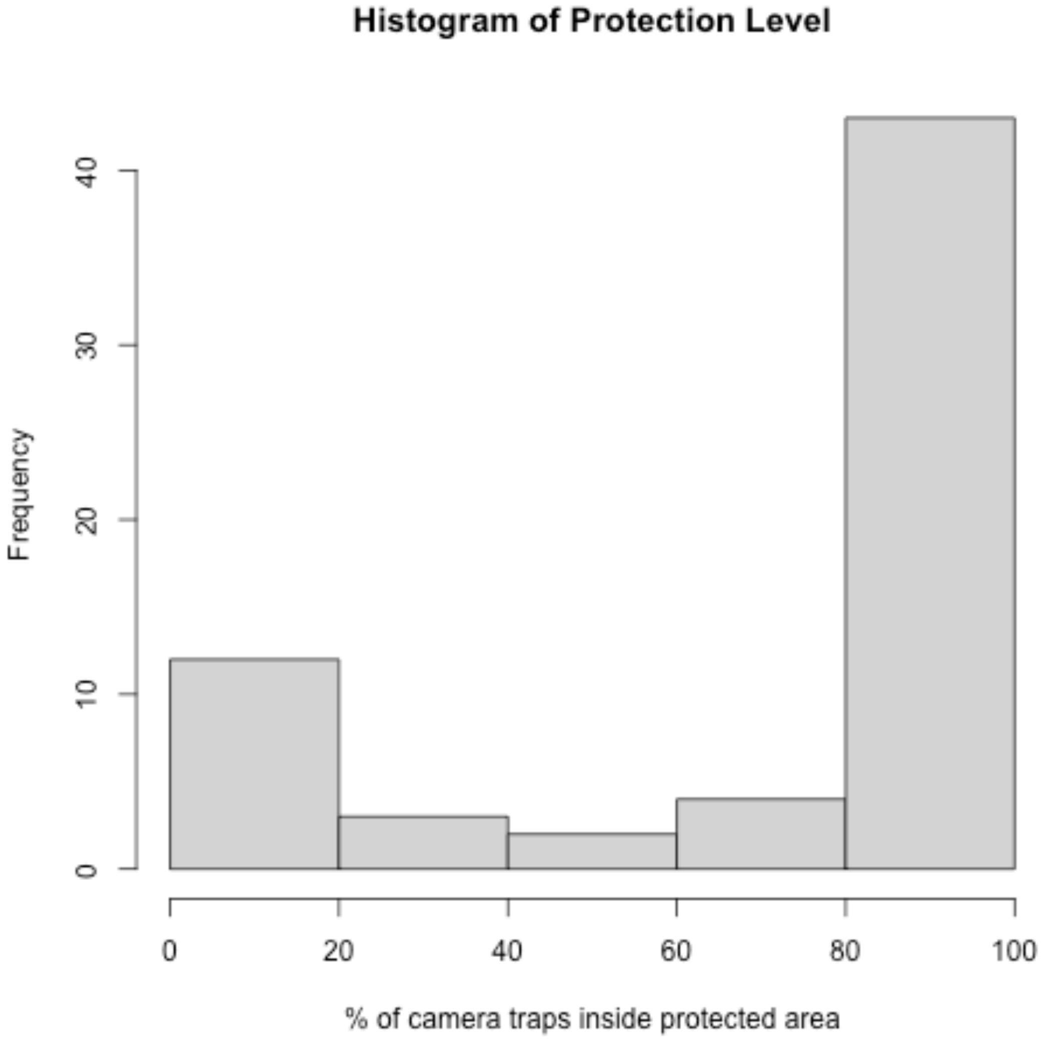


**Figure S3.** The percentage of camera traps at our study sites that are located inside a protected area.

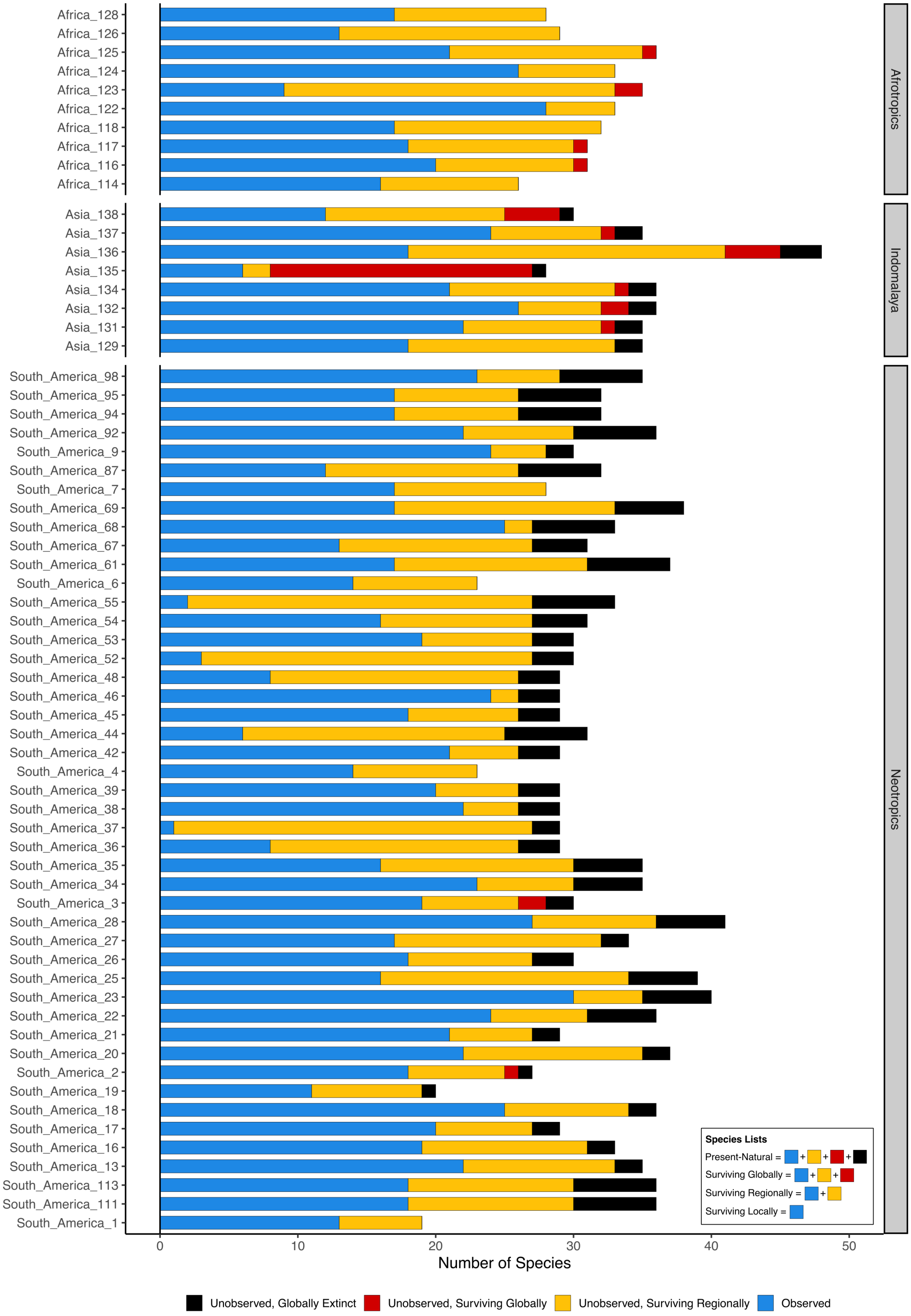


**Figure S4.** The Species Lists for each site.

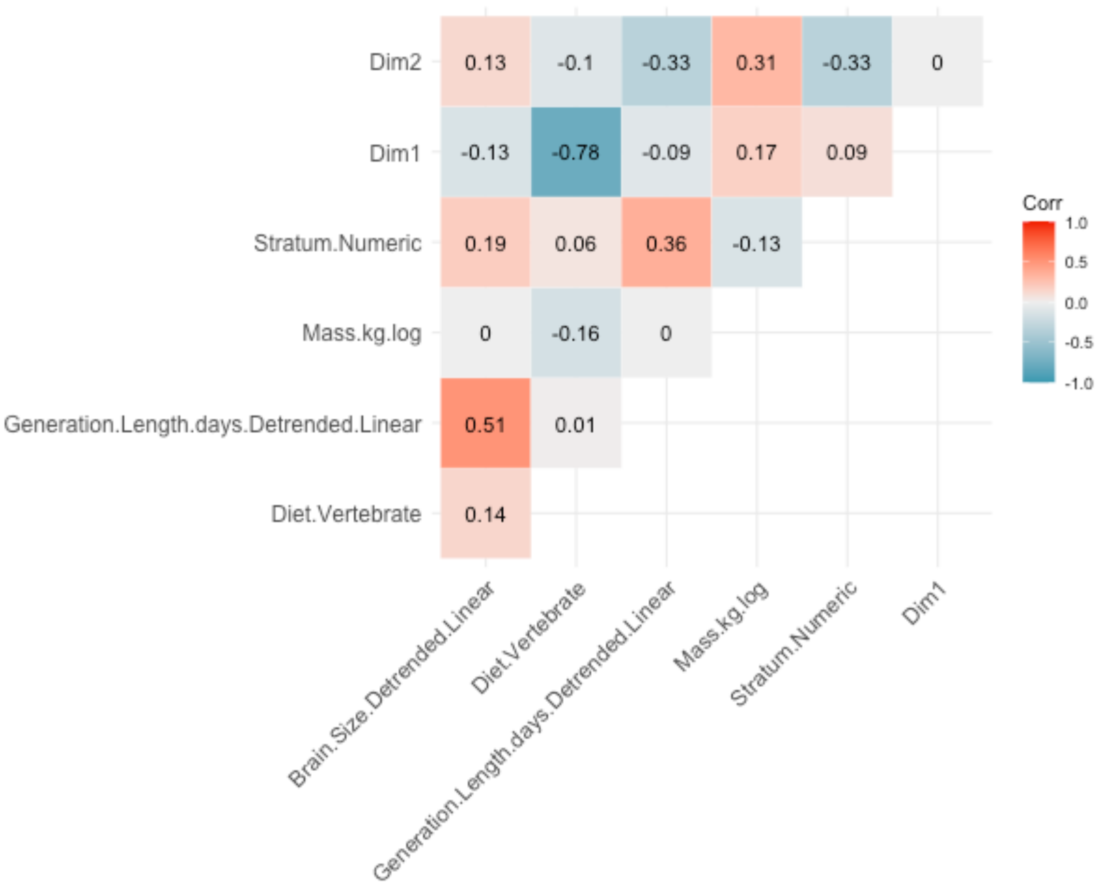


**Figure S5:** A correlation matrix showing the correlations between all trait and phylogenetic variables used in the analysis.

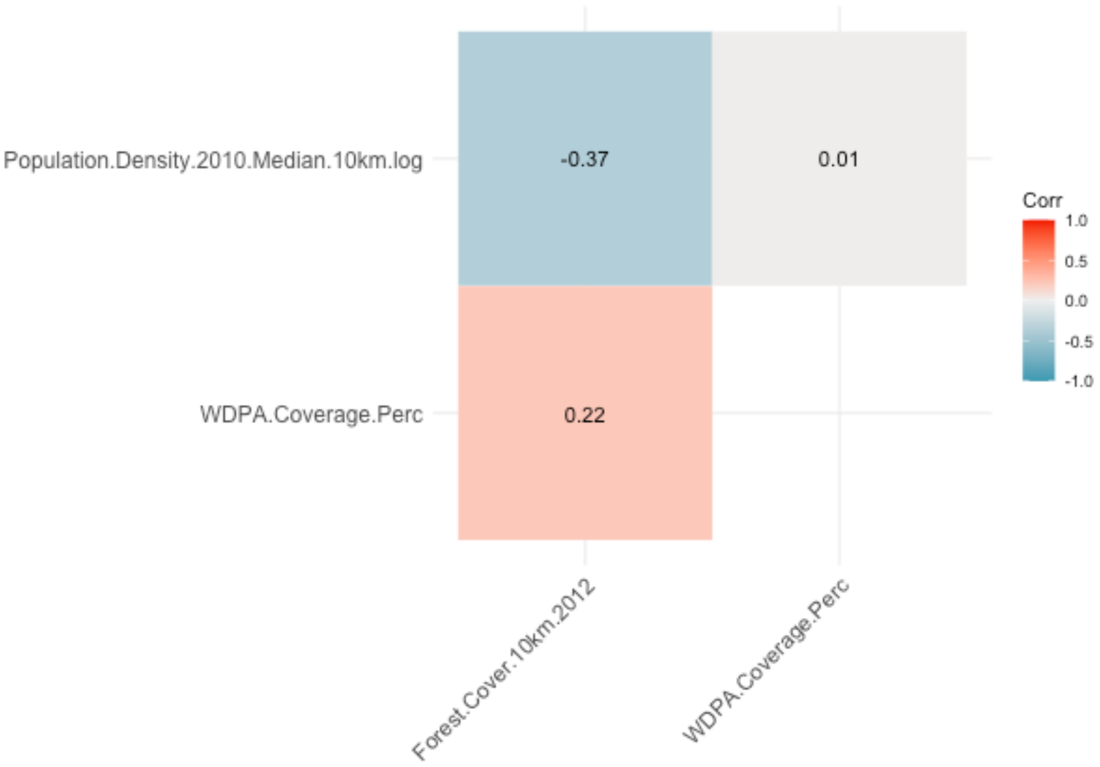


**Figure S6.** *A correlation matrix showing the correlations between all site variables used in the analysis.*

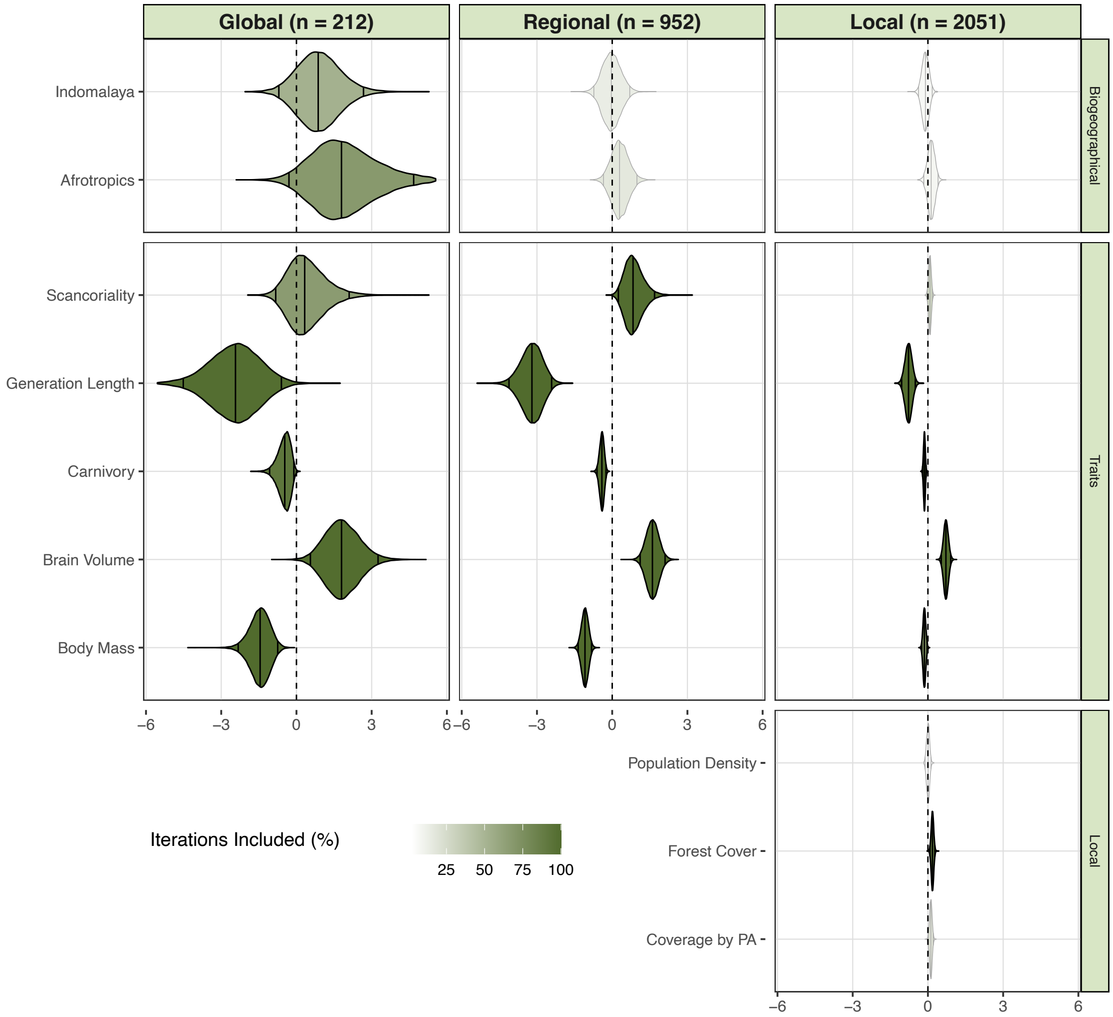


**Figure S7. Posterior distributions of the coefficient estimates of spatial-only model, using a 5 km buffer to extract site data.** Vertical lines inside the violin plots show the 95% credible Intervals and the median values. The colour intensity of the plots show how often each variable was included in the model. Variables included in more than 50% of the iterations were drawn with black borders. Variables included in fewer than 50% of the iterations have been drawn using gray borders. The Neotropics are the intercept level to which the other realms are compared.

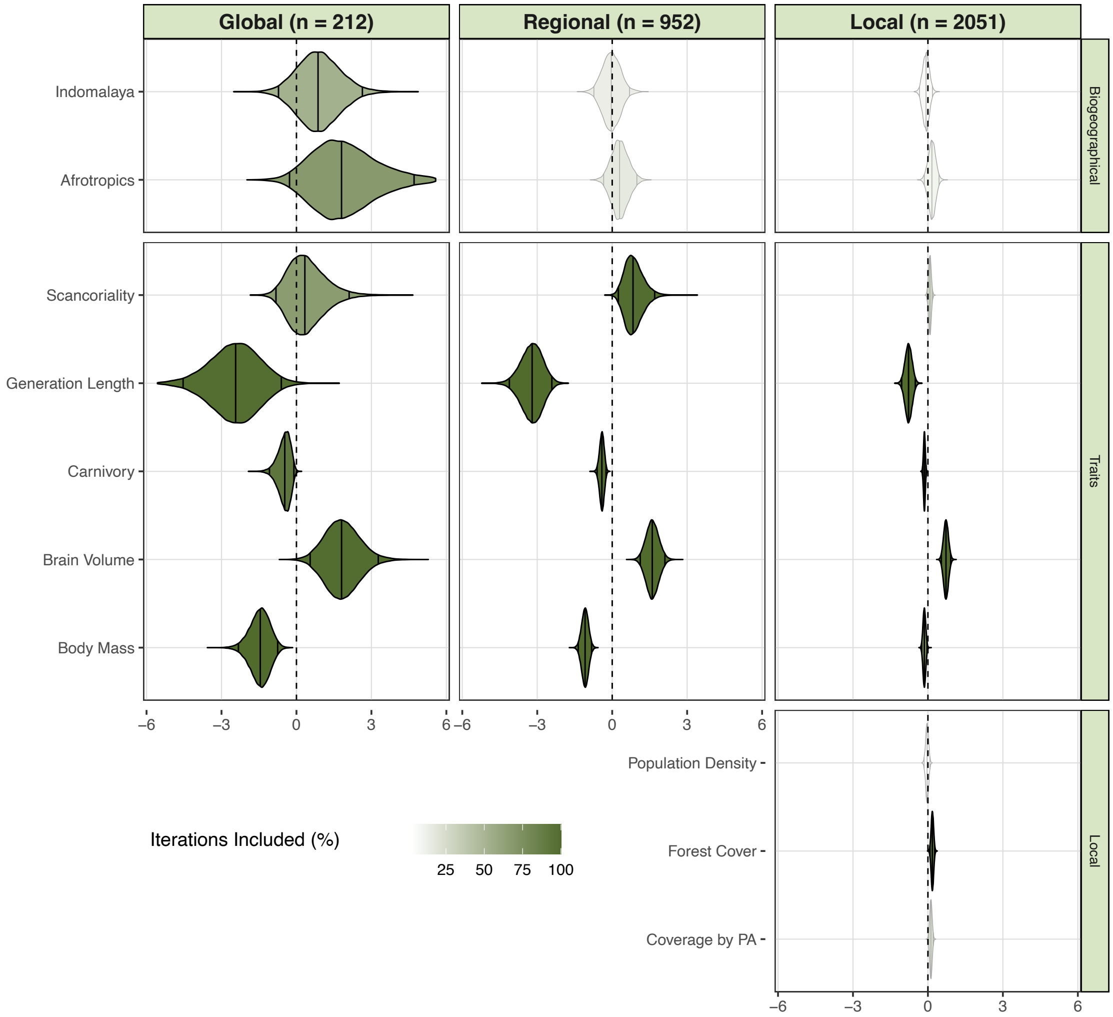


**Figure S8. Posterior distributions of the coefficient estimates of spatial-only model, using a 20 km buffer to extract site data**. Vertical lines inside the violin plots show the 95% credible Intervals and the median values. The colour intensity of the plots show how often each variable was included in the model. Variables included in more than 50% of the iterations were drawn with black borders. Variables included in fewer than 50% of the iterations have been drawn using gray borders. The Neotropics are the intercept level to which the other realms are compared.

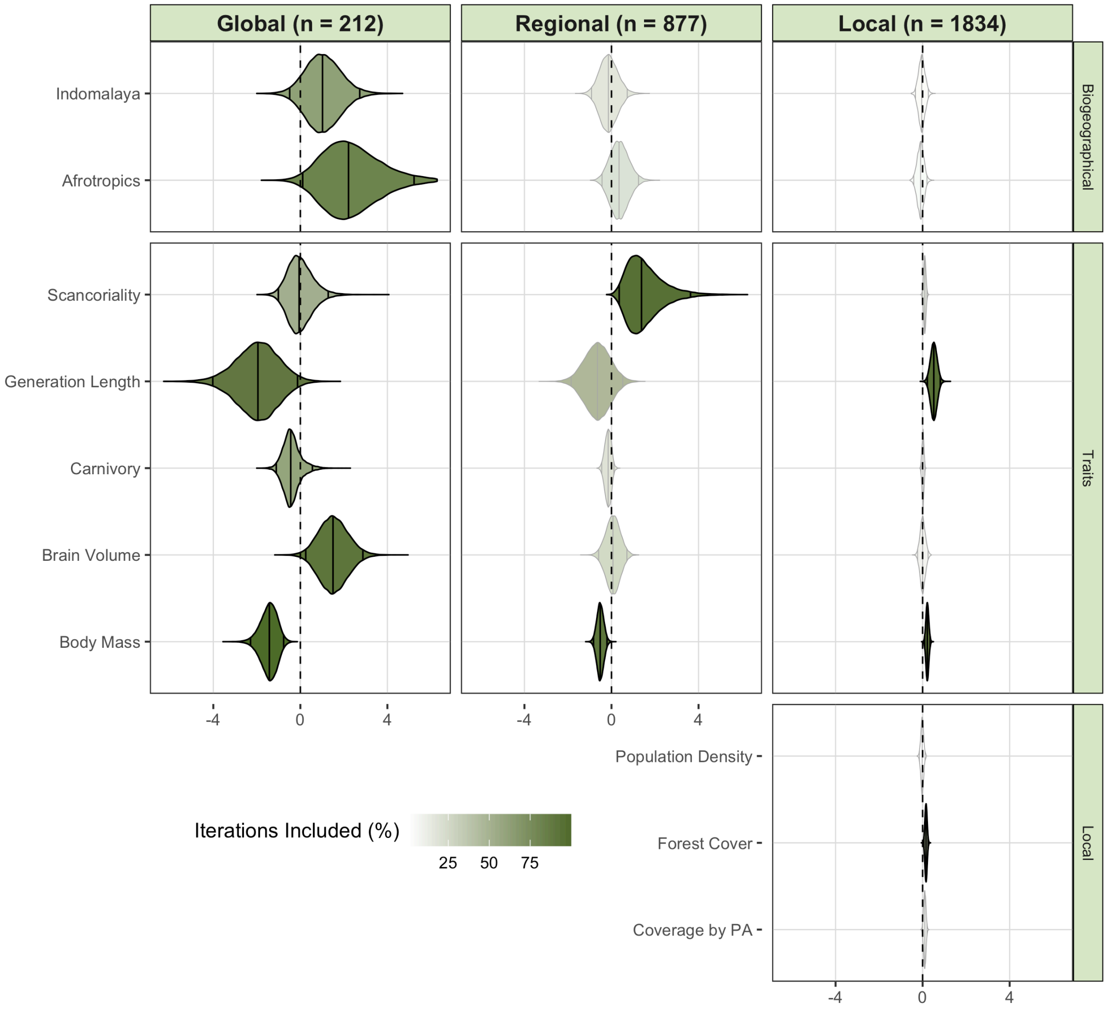


**Figure S9. Posterior distributions of the coefficient estimates of spatio-temporal model, using a 5 km buffer to extract site data**. Vertical lines inside the violin plots show the 95% credible Intervals and the median values. The colour intensity of the plots show how often each variable was included in the model. Variables included in more than 50% of the iterations were drawn with black borders. Variables included in fewer than 50% of the iterations have been drawn using gray borders. The Neotropics are the intercept level to which the other realms are compared.

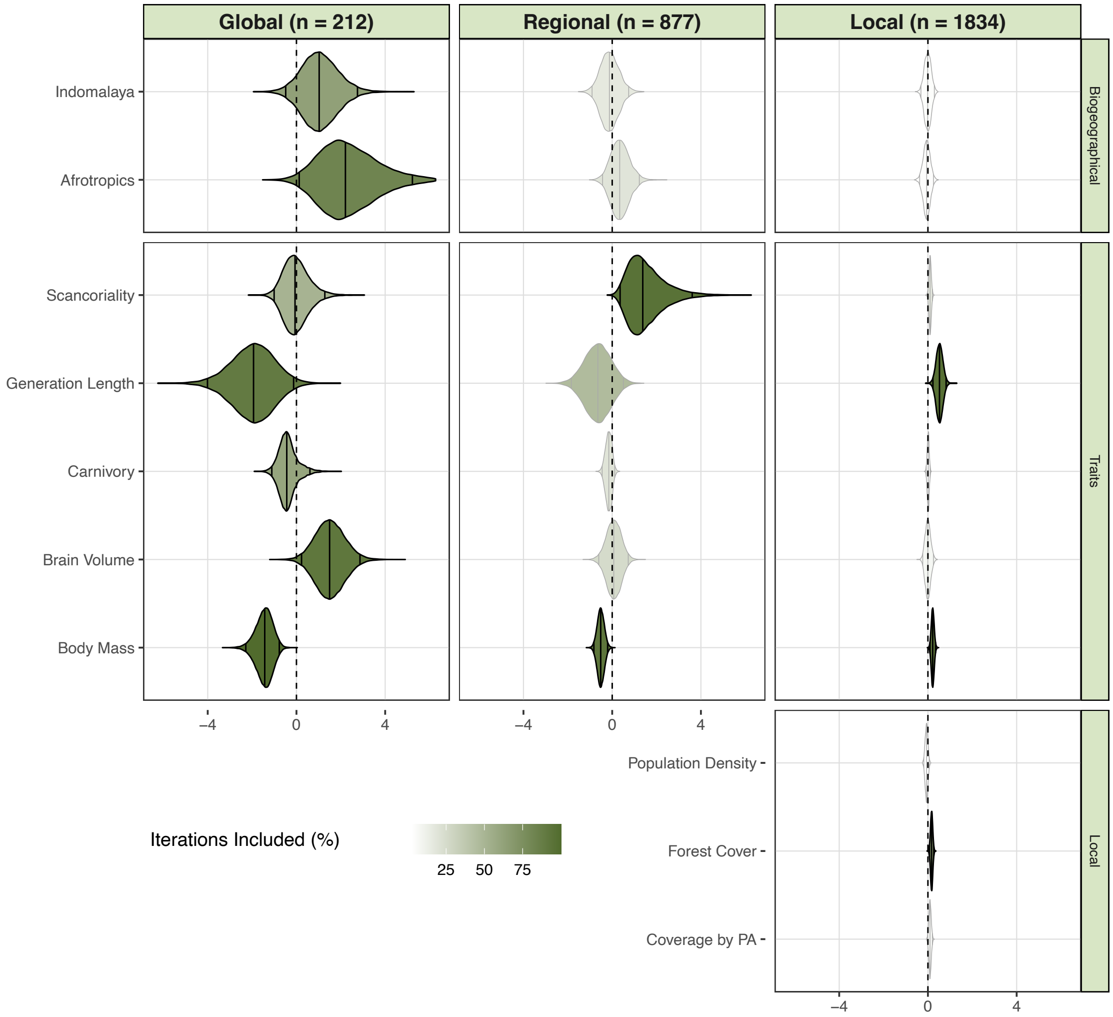


**Figure S10. Posterior distributions of the coefficient estimates of spatial-temporal model, using a 20 km buffer to extract site data.** Vertical lines inside the violin plots show the 95% credible Intervals and the median values. The colour intensity of the plots show how often each variable was included in the model. Variables included in more than 50% of the iterations were drawn with black borders. Variables included in fewer than 50% of the iterations have been drawn using gray borders. The Neotropics are the intercept level to which the other realms are compared.

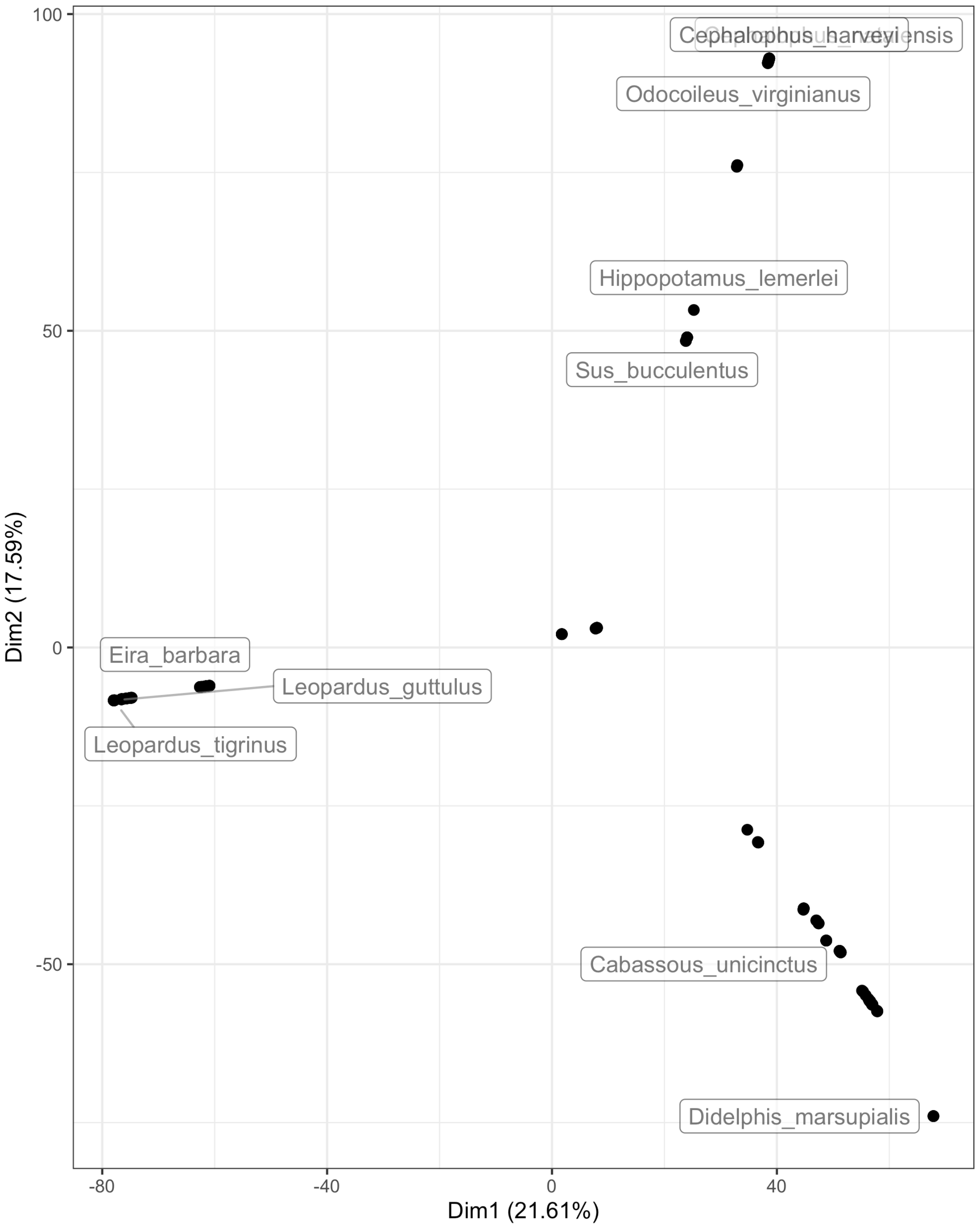


**Figure S11. The Phylogenetic Eigenvectors showing the phylogenetic relationships between the species in our analysis.** Species with high values on phylogenetic dimension 1 tended to be marsupials or rodents (e.g. Didelphis, Dasyprocta). Species with low values tended to be carnivorans (e.g. Leopardus, Prionailurus, Puma). Species with high values along dimension 2 tended to be ungulates (e.g. Cephalophus, Mazama).

**Table S1. Global predictors of the spatial-only model** - the median estimate, Bayesian 95% credible interval and inclusion probability of the selected global scale coefficients.

| Name | Median | CI.2.5 | CI.97.5 | Inclusion Probability |
|---|---|---|---|---|
| Indomalaya | 0.86 | -0.72 | 2.68 | 0.52 |
| Afrotropics | 1.83 | -0.29 | 5.30 | 0.68 |
| Brain Volume | 1.81 | 0.55 | 3.27 | 0.99 |
| Vertebrate Carnivory | -0.47 | -1.08 | -0.11 | 0.93 |
| Generation Length | -2.43 | -4.57 | -0.62 | 0.98 |
| Body Mass | -1.45 | -2.33 | -0.76 | 1.00 |
| Scansoriality | 0.34 | -0.81 | 2.09 | 0.66 |

**Table S2. Regional predictors of the spatial-only model** - the median estimate, Bayesian 95% credible interval and inclusion probability of the selected regional scale coefficients.

| Name | Median | CI.2.5 | CI.97.5 | Inclusion Probability |
|---|---|---|---|---|
| Indomalaya | -0.04 | -0.72 | 0.65 | 0.14 |
| Afrotropics | 0.30 | -0.34 | 0.97 | 0.17 |
| Brain Volume | 1.61 | 1.13 | 2.10 | 1.00 |
| Vertebrate Carnivory | -0.41 | -0.60 | -0.25 | 1.00 |
| Generation Length | -3.20 | -4.11 | -2.42 | 1.00 |
| Body Mass | -1.09 | -1.36 | -0.82 | 1.00 |
| Scansoriality | 0.84 | 0.25 | 1.73 | 0.99 |

**Table S3. Local predictors of the spatial-only model** - the median estimate, Bayesian 95% credible interval and inclusion probability of the selected local scale coefficients.

| Name | Median | CI.2.5 | CI.97.5 | Inclusion Probability |
|---|---|---|---|---|
| Indomalaya | -0.07 | -0.33 | 0.19 | 0.06 |
| Afrotropics | 0.16 | -0.14 | 0.43 | 0.10 |
| Brain Volume | 0.72 | 0.54 | 0.91 | 1.00 |
| Vertebrate Carnivory | -0.14 | -0.21 | -0.08 | 1.00 |
| Generation Length | -0.77 | -1.05 | -0.50 | 1.00 |
| Body Mass | -0.14 | -0.25 | -0.04 | 0.77 |
| Scansoriality | 0.08 | -0.03 | 0.18 | 0.30 |
| Forest.Cover.20km.2012 | 0.18 | 0.08 | 0.27 | 0.90 |
| Protected Area Coverage | 0.12 | 0.02 | 0.22 | 0.26 |
| Population.Density.2010.Median.20km.log | -0.04 | -0.15 | 0.07 | 0.03 |

**Table S4. Global predictors of the spatio-temporal model** - the median estimate, Bayesian 95% credible interval and inclusion probability of the selected global scale coefficients.

| Name | Median | CI.2.5 | CI.97.5 | Inclusion Probability |
|---|---|---|---|---|
| Indomalaya | 1.03 | -0.45 | 2.74 | 0.63 |
| Afrotropics | 2.24 | 0.17 | 5.75 | 0.83 |
| Brain Volume | 1.50 | 0.26 | 2.87 | 0.92 |
| Vertebrate Carnivory | -0.43 | -1.10 | 0.58 | 0.59 |
| Generation Length | -1.93 | -3.98 | -0.13 | 0.90 |
| Body Mass | -1.42 | -2.28 | -0.76 | 1.00 |
| Scansoriality | -0.06 | -0.98 | 1.30 | 0.51 |

**Table S5. Regional predictors of the spatio-temporal model** - the median estimate, Bayesian 95% credible interval and inclusion probability of the selected regional scale coefficients.

| Name | Median | CI.2.5 | CI.97.5 | Inclusion Probability |
|---|---|---|---|---|
| Indomalaya | -0.14 | -0.89 | 0.73 | 0.17 |
| Afrotropics | 0.35 | -0.43 | 1.24 | 0.19 |
| Brain Volume | 0.07 | -0.60 | 0.70 | 0.25 |
| Vertebrate Carnivory | -0.15 | -0.44 | 0.12 | 0.19 |
| Generation Length | -0.64 | -1.83 | 0.50 | 0.45 |
| Body Mass | -0.52 | -0.82 | -0.20 | 0.93 |
| Scansoriality | 1.39 | 0.38 | 3.65 | 0.95 |

**Table S6. Local predictors of the spatio-temporal model** - the median estimate, Bayesian 95% credible interval and inclusion probability of the selected local scale coefficients.

| Name | Median | CI.2.5 | CI.97.5 | Inclusion Probability |
|---|---|---|---|---|
| Indomalaya | -0.02 | -0.31 | 0.27 | 0.04 |
| Afrotropics | -0.05 | -0.36 | 0.23 | 0.04 |
| Brain Volume | 0.00 | -0.23 | 0.24 | 0.07 |
| Vertebrate Carnivory | 0.01 | -0.07 | 0.10 | 0.04 |
| Generation Length | 0.52 | 0.23 | 0.82 | 0.96 |
| Body Mass | 0.22 | 0.10 | 0.34 | 0.92 |
| Scansoriality | 0.10 | -0.01 | 0.20 | 0.16 |
| Forest.Cover.10km.2012 | 0.17 | 0.07 | 0.26 | 0.70 |
| Protected Area Coverage | 0.09 | -0.01 | 0.20 | 0.06 |
| Population.Density.2010.Median.10km.log | -0.05 | -0.17 | 0.07 | 0.03 |

**Table S7. Trait Data Completion** - Species traits and the degree of data completion before imputation (in %) for all species (n = 210), globally extinct species (n = 11), and extant species (n = 199).

| Variable | All.Species | Extinct.Only | Extant.Only |
|---|---|---|---|
| Diet | 96.67 | 72.73 | 97.99 |
| Brain Volume | 62.86 | 27.27 | 64.82 |
| Generation Length | 75.71 | 9.09 | 79.40 |
| Body Mass | 98.10 | 100.00 | 97.99 |
| Scansoriality | 100.00 | 100.00 | 100.00 |

**Vignette S1. (separate file)**

*Vignette.html*– An html vignette explaining how the model works.

**Data S1 (separate file)**

*SI_Data_1_rarefaction_curves.pdf* – A .pdf file showing the species accumulation curves of each site.

**Data S2 (separate file)**

*SI_Data_2_excluded_species.csv* – A .csv file listing the species that we excluded from the analysis.

**Data S3 (separate file)**

*SI_Data_3_range_inconsistencies.csv* – A .csv file reporting the inconsistencies between present-natural range maps and the camera trap data.

**Data S4. (separate file)**

*SI_Data_4_Trait_Array.rds* – An .rds file containing the species trait data including imputed values, organised as an array.

**Data S5 (separate file)**

*SI_Data_5_Species_Traits_and_Metadata.csv* – A .csv file with trait data and metadata, but without imputed values.

**Data S6 (separate file)**

*SI_Data_6_site_info.csv* – A .csv file containing site data.

**Data S7 (separate file)**

*SI_Data_7_species_park_list.csv* – A .csv file containing the species presence-absences
