## Supplementary material for "Predictors of extinction risk in large tropical forest mammals: from global to local": Vignette.html: Vignette.html

Vignette Extinction Scales


#### Table of contents

- Prepare Script
  - Load R packages
  - Load the Data
  - Model Specifications
- Prepare the Data
  - Prepare Presence-Absence Data
  - Prepare Trait Data
    - Inspect Trait Distributions
  - Prepare Site Data
  - Select Model Predictors
  - Standardise Data
  - Merge the Biogeographical and Site Data
  - Trait Lookup Table
  - Species X Site lookup tables
- Prepare the Model
  - Define Nimble Model
- Define Model Inputs
  - Define the Data
  - Define the Constants
  - Define Initial Values
  - Specify the parameters to retrieve
- Fit the Model
- Process Nimble Output
  - Show Traceplots
  - Plot Coefficients

### Vignette Extinction Scales

Author

Simon D. Schowanek, Pierre Dupont and Richard Bischof

#### 2024-08-19

This script is part of the supplementary materials of the manuscript: “Predictors of extinction risk in large tropical forest mammals: from global to local”. It illustrates how the models used in the manuscript work. To make changes to the model, go to the section “Model Specifications”.

### Prepare Script

#### Load R packages

```
library(coda)                      # Import the MCMC diagnostic tools
library(nimble)                    # Import the NIMBLE subroutines
library(ggplot2)
library(dplyr)
library(reshape2)
library(ggridges)
library(RColorBrewer)
```

#### Load the Data

Here, we load the data and prepare it for the analysis.

```
## – LOAD DATA ––––
#–––––––––––––––––––––––––––––––––––––––––––––––––

## Load ocurrence Data (and rename biogeographic regions to match names in the paper)
df.presence <- read.csv("./Vignette_Submission_files/data/SI_Data_7_species_park_list.csv") %>%
  mutate( Bio.Realm = recode(Bio.Region,
                              'Africa' = 'Afrotropics',
                              'Asia' = 'Indomalaya',
                              'South_America' = 'Neotropics'))


## Load Site data (and filter out unused sites)
df.sites <- read.csv("./Vignette_Submission_files/data/SI_Data_6_site_info.csv") %>%
  filter(., Polygon.ID %in% unique(df.presence$Polygon.ID))  


## Load Species Traits data
traits.ar <- readRDS("./Vignette_Submission_files/data/SI_Data_4_Trait_Array.rds")
```

The trait data also contains the phylogenetic eigenvectors that quantify the phylogenetic relationships between species. We ran PCoAs on the phylogenetic distances between species (see the paper for more details). The trait dataset contains the coordinates of those PCoAs. We can use these coordinates to plot the relatedness between species and to account for this phylogenetic relatedness in our model. However, each species has a range of coordinates and we need to simplify these data. The following section takes the PCoA coordinates of the Phylogenetic Eigenvectors and replaces them with the mean value. We do this because the model has convergence issues when we give it the full range of possible values.

```
for(i.sp in 1:dim(traits.ar)[2]){
  traits.ar[ ,i.sp,"Dim1"] = mean(traits.ar[ ,i.sp,"Dim1"])
  traits.ar[ ,i.sp,"Dim2"] = mean(traits.ar[ ,i.sp,"Dim2"])
  traits.ar[ ,i.sp,"Dim3"] = mean(traits.ar[ ,i.sp,"Dim3"])
}
```

Finally, a few things need to be defined

```
## Name The Three Scales
scale.names = c("Global", "Regional", "Local")

## Number of Species
n.species = length(unique(df.presence$Binomial))

## Species List
v.species.list = sort(unique(df.presence$Binomial))
```

#### Model Specifications

If you want to test different version of the model, this is where you should do it.

###### Spatial-Only or Spatio-Temporal Model?

Should we run the “Spatial-Only” model (Spatial.Only = T) or the “Spatio-Temporal” model (Spatial.Only = F)?

```
Spatial.Only = T
```

###### Phylogenetic Correction

Should the model run with or without a phylogenetic correction? Note that turning on the Phylogenetic correction takes more time.

```
Phylogenetic.Correction = F
```

###### Bayesian Model Specifications

Here, we specify the details of the Bayesian Model.

```
## MODEL SPECS
df.model.specs = cbind.data.frame( mcmc = "cMCMC",
                                   nburnin = 10 * 10^0,
                                   niter= 3 * 10^3, 
                                   nchains = 3,
                                   thin = 1,
                                   k.Inclusion.Treshold = 0.05)
```

###### Buffer Size for Local Variables

In the original model, the local site variables were extracted using a 10 km buffer. It is possible to also run the model using a 5 or a 20 km buffer.

```
buffer.radius = 10 # 5, 10, or 20 km?
df.model.specs$buffer.radius = buffer.radius
```

### Prepare the Data

#### Prepare Presence-Absence Data

Here, we prepare the presence-absence data for the model. Before, we do so we will visualise what the data look like. We will create a figure that consists of four columns representing the Present-natural, Surviving Globally, Surviving regionally, and Surviving Locally species respectively. Each row represents a species x site combination (e.g. *Panthera tigris* occurring in site Asia\_1). Red means extant. Yellow means extinct. Note, that species can only occur at a given scale if they exist at the scale above it (e.g. a species can not be locally present if they are globally extinct)

```
## Select Columns
df.pres.abs <- df.presence[ ,c("Present.Natural",
                              "Present.Globally",
                              "Present.Regionally",
                              "Present.Locally")]

## Show nested Structure of extinctions
image(t(df.pres.abs))
```

Next, we will create the actual datasets that will be used in the model. We start with the global scale dataset.

```
## CREATE IDENTIFIERS
df.presence$ID.Global = paste0(df.presence$Binomial, "_", df.presence$Bio.Realm)
df.presence$ID.Regional = paste0(df.presence$Binomial, "_", df.presence$BRU)
df.presence$ID.Local = paste0(df.presence$Binomial, "_", df.presence$Polygon.ID)


##-- GLOBAL

## Select Columns
df.pres.global <- unique(df.presence[ ,c("Binomial",
                                        "Bio.Realm",
                                        "ID.Global",
                                        "Present.Globally")])

## Add Rownames
rownames(df.pres.global) = df.pres.global$ID.Global
```

From here on, the process is slightly different for the spatial-only and the spatio-temporal model, because the model type changes which species are included in the regional and the local levels. When the spatio-temporal is run, all globally extinct species are excluded from the species pool. The following code is similar to the code for the global level but it adds an if statement to account for the species removal in the spatio-temporal model.

```
##-- REGIONAL

## Identify Globally Extant and Extinct Species
v.extant.global <- df.pres.global[df.pres.global$Present.Globally == 1,
                                 "Binomial"]

## Select Columns
df.pres.regional <- unique(df.presence[ ,c("Binomial",
                                           "Bio.Realm",
                                           "ID.Global",
                                           "ID.Regional",
                                           "BRU",
                                           "Present.Regionally")])

## Add Rownames
rownames(df.pres.regional) = df.pres.regional$ID.Regional 

## Remove species that are globally extinct (WHEN RUNNING SPATIO-TEMPORAL MODEL)
if(Spatial.Only == F){
  df.pres.regional <- df.pres.regional %>%
    filter(Binomial %in% v.extant.global) 
} # close if
```

In the same vein, the code for the local scale is similar to the code for the regional scale, but it adds an if statement to account for the species removal in the spatio-temporal model.

```
#- LOCAL

## Check which species are regionally extant
v.extant.regional <- df.pres.regional[df.pres.regional$Present.Regionally == 1,
                                      "ID.Regional"]

## Select Columns
df.pres.local <- unique(df.presence[ ,c("Binomial",
                                       "Bio.Realm",
                                       "ID.Regional",
                                       "Polygon.ID",
                                       "Present.Locally")])

## Add Rownames
rownames(df.pres.local) = paste0(df.pres.local$Binomial, "_", df.pres.local$Polygon.ID)

## Remove Species that are regionally extinct (WHEN RUNNING SPATIO-TEMPORAL MODEL)
if(Spatial.Only == F){
df.pres.local <- df.pres.local %>% filter(ID.Regional %in% v.extant.regional) 
} # close i
```

Finally, we remove redundant columns.

```
df.pres.global <- df.pres.global %>%
  select(Binomial, Bio.Realm, Present.Globally)

df.pres.regional <- df.pres.regional %>%
  select(Binomial, Bio.Realm, BRU, Present.Regionally)

df.pres.local <- df.pres.local %>%
    select(Binomial, Bio.Realm, Polygon.ID, Present.Locally)
```

#### Prepare Trait Data

This code takes the trait data and calculates the means and standard deviations. These will later be used in the model.

```
## TURN ARRAY INTO DF
df.traits.mean<-apply(traits.ar,c(2,3),mean,na.rm=TRUE)
```

The trait data contains known values, as well as uncertain values (values with an imputed distribution for the trait). Therefore, we will specify standard deviations around the mean values. Species with known traits will have their standard deviations set to NA, to reflect that there is no uncertainty. Species with uncertain traits will receive a normal standard deviation value.

```
## – TURN ARRAY INTO DF WITH SD VALUES ––––
#–––––––––––––––––––––––––––––––––––––––––––––––––

## Create SD array
df.traits.sd = apply(traits.ar, c(2:3), sd)

## Turn species with SD == 0 TO NA
df.traits.sd[(df.traits.sd == 0)] = NA

## Show how many uncertain values there are for every trait
apply(df.traits.sd, 
      2,
      function(x){1-length(which(is.na(x)))/nrow(df.traits.sd)})
```

```
                                 Mass.g                              Diet.Plant 
                             0.01843318                              0.03686636 
                        Diet.Vertebrate              Generation.Length.days.log 
                             0.03686636                              0.25345622 
        Endocast.Volume.ml.combined.log                         Stratum.Numeric 
                             0.36866359                              0.00000000 
                                   Dim1                                    Dim2 
                             0.00000000                              0.00000000 
                                   Dim3                  Generation.Length.days 
                             0.00000000                              0.25345622 
            Endocast.Volume.ml.combined                     Relative.Brain.Size 
                             0.36866359                              0.36866359 
                Relative.Brain.Size.log                             Mass.kg.log 
                             0.36866359                              0.01843318 
            Brain.Size.Detrended.Linear Generation.Length.days.Detrended.Linear 
                             1.00000000                              1.00000000
```

##### Inspect Trait Distributions

This code randomly selects 30 species and plots their trait distribution. Some traits only have one unique value (if traits are known). They show up in the plots as perfect normal distributions. Species with uncertain trait values will have more irregular and more spread out distributions. For visualisation purposes, we show the first 3 plots.

```
idx <- sample(dim(traits.ar)[2],min(dim(traits.ar)[2],30))#---draw a random sample of species

plot.list.1 <- lapply(1:dim(traits.ar)[3],function(i.cov){
   
  temp <- as.data.frame(traits.ar[ ,idx,i.cov])
  temp <- melt(temp) 
  
  ggplot() + 
    geom_density_ridges(data=temp,aes(x=value,
                                      y=variable,
                                      fill=variable, 
                                      color=variable),
                        alpha=0.25,
                        bandwidth = 0.01) +
    ggtitle(dimnames(traits.ar)[[3]][i.cov]) + 
    theme(legend.position = "none")
})

## Plot Trait Distributions
plot.list.1[1:3]
```

```
[[1]]
```

```
[[2]]
```

```
[[3]]
```

#### Prepare Site Data

```
#-- Create Table with Site Location Info
df.site.info = unique(df.presence[,c("Bio.Realm",
                                     "BRU",
                                     "Polygon.ID")])

#-- merge site info data with 
df.sites = merge(df.sites, df.site.info, by = "Polygon.ID")

#-- Add Rownames
rownames(df.sites) = df.sites$Polygon.ID
```

#### Select Model Predictors

```
##– TRAITS ––––
#–––––––––––––––––––––––––––––––––––––––––––––––––

## SELECT TRAIT VARIABLES
v.trait.cov = sort(c("Mass.kg.log",
                     "Diet.Vertebrate",
                     "Stratum.Numeric",
                     "Generation.Length.days.Detrended.Linear",
                     "Brain.Size.Detrended.Linear"
                     )) # select intrinsic variables


## ADD PHYLOGENETIC CORRECTION TO TRAIT MATRIX (IF PHYLOGENETIC CORRECTION IS TURNED ON)
if(Phylogenetic.Correction == T){
  v.trait.cov = c(v.trait.cov, "Dim1" ,"Dim2")
}


##– OTHER VARIABLES ––––
#–––––––––––––––––––––––––––––––––––––––––––––––––

## Name biogeographical realms
v.ext.cov.shared = c("Indomalaya", "Afrotropics")

## Select Site Variables
v.ext.cov.local = c(paste0("Population.Density.2010.Median.", buffer.radius, "km.log"),
                    paste0("Forest.Cover.", buffer.radius, "km.2012"),
                    "WDPA.Coverage.Perc"
                    ) # Select External Variables

v.ext.cov = c(v.ext.cov.shared, v.ext.cov.local) # combine


##– COMBINED VARIABLE LIST ––––
#–––––––––––––––––––––––––––––––––––––––––––––––––
v.selected.predictors.global <- c(v.ext.cov.shared,
                                  v.trait.cov) # Combine predictors into one variable

v.selected.predictors.regional <- c(v.ext.cov.shared,
                                   v.trait.cov) # Combine predictors into one variable

v.selected.predictors.local <- c(v.ext.cov,
                                 v.trait.cov) # Combine predictors into one variable

v.selected.predictors <- v.selected.predictors.local # this is the total set of predictors
```

#### Standardise Data

```
## – STANDARDISE THE TRAIT MEANS ––––
#–––––––––––––––––––––––––––––––––––––––––––––––––

## Standardise Trait Means
df.traits.mean.scaled = df.traits.mean[ ,colnames(df.traits.mean) %in% v.selected.predictors] 

## select variables
df.traits.mean.scaled = df.traits.mean.scaled[ ,v.trait.cov] # reorder variables

## scale variables
df.traits.mean.scaled = apply(df.traits.mean.scaled,
                              2,
                              FUN = function(a){(a - mean(a)) / sd(a)})
```

We have standardised the predictor variables, but to make sure we include the trait uncertainty in the model, we also need to include the original trait array in the model (as this contains the estimated distributions for each trait). To make sure these data are comparable we standardise them as well.

```
#–– STANDARDISE THE TRAIT ARRAY
#–––––––––––––––––––––––––––––––––––––––––––––––––

## PREPARE DATA FOR STANDARDISATION
ar.traits.scaled = traits.ar[,,v.trait.cov]
v.colnames = dimnames(ar.traits.scaled)[[3]] # update colnames

## STANDARDISE VALUES IN TRAIT ARRAY
for (i.col in 1:dim(ar.traits.scaled)[3]) {
  
  colname = v.colnames[i.col] # select column
  
  ar.traits.scaled[ , ,colname] = 
    (ar.traits.scaled[ , ,colname] - 
       mean(df.traits.mean.scaled[,colname])) / sd(df.traits.mean.scaled[,colname])
} # close for loop
```

Because Stratum Numeric is a binary variable we do not scale it. We overwrite it again.

```
ar.traits.scaled[ , ,"Stratum.Numeric"] = traits.ar[ , ,"Stratum.Numeric"]
```

The trait data contains known values, as well as uncertain values (values with an imputed distribution for the trait). Therefore, we will specify standard deviations around the mean values. Species with known traits will have their standard deviations set to NA, to reflect that there is no uncertainty. Species with uncertain traits will receive a normal standard deviation value.

```
#–– TURN ARRAY INTO DF WITH SD VALUES 
#–––––––––––––––––––––––––––––––––––––––––––––––––

## Create SD array
df.traits.sd.scaled = apply(ar.traits.scaled, c(2:3), sd)

## Turn species with SD == 0 TO NA
df.traits.sd.scaled[(df.traits.sd.scaled == 0)] = NA

## Show how many uncertain values there are for every trait
apply(df.traits.sd.scaled,
      2,
      function(x){1-length(which(is.na(x)))/nrow(df.traits.sd.scaled)})
```

```
            Brain.Size.Detrended.Linear                         Diet.Vertebrate 
                             1.00000000                              0.03686636 
Generation.Length.days.Detrended.Linear                             Mass.kg.log 
                             1.00000000                              0.01843318 
                        Stratum.Numeric 
                             0.00000000
```

```
#–– STANDARDISE SITE DATA ––––
#–––––––––––––––––––––––––––––––––––––––––––––––––
df.sites.scaled =
  df.sites[ ,colnames(df.sites) %in% v.selected.predictors] # create new dataset

df.sites.scaled = apply(df.sites.scaled,
                        2,
                        FUN = function(a){(a - mean(a)) / sd(a)})
```

#### Merge the Biogeographical and Site Data

```
##– CHANGE FORMAT OF THE BIOGEOGRAPHICAL DATA 
#–––––––––––––––––––––––––––––––––––––––––––––––––

#- CREATE LIST
l.site.data = list(unique(df.pres.global[,c( "Binomial", "Bio.Realm")]),
                   unique(df.pres.regional[,c( "Binomial","Bio.Realm", "BRU")]),
                   unique(df.pres.local[,c( "Binomial","Bio.Realm", "Polygon.ID")]))
names(l.site.data) = scale.names

## FILL LIST
l.biogeo <- lapply(l.site.data, function(x){
  x[, "Indomalaya"] = 0
  x[, "Afrotropics"] = 0
  x[which(x[,"Bio.Realm"] == "Indomalaya"), "Indomalaya"] = 1
  x[which(x[,"Bio.Realm"] == "Afrotropics"), "Afrotropics"] = 1
  x
}) # close apply


##– CHANGE FORMAT OF THE BIOGEOGRAPHICAL DATA ––––
#–––––––––––––––––––––––––––––––––––––––––––––––––
df.sites.scaled = data.frame(df.sites.scaled) # turn matrix)array into df
df.sites.scaled[,"Polygon.ID"] = rownames(df.sites.scaled) # turn rownames into column (for merge later)

l.site.data$Global <-
  l.biogeo$Global[,colnames(l.biogeo$Global) %in% v.ext.cov.shared] # select relevant variables

l.site.data$Regional <-
  l.biogeo$Regional[,colnames(l.biogeo$Regional) %in% v.ext.cov.shared] # select relevant variables

## LOCAL
l.biogeo$Local <-  
  l.biogeo$Local[,colnames(l.biogeo$Local) %in% c("Polygon.ID",
                                                  "Indomalaya",
                                                  "Afrotropics")] # select relevant variables

l.site.data$Local <- 
  merge(l.biogeo$Local, df.sites.scaled, by = "Polygon.ID", all.x = T)  # merge

l.site.data$Local <-  
  l.site.data$Local[,colnames(l.site.data$Local) %in% v.selected.predictors] # select only numeric predictors
```

#### Trait Lookup Table

Some of the trait values in our dataset are known, others are unknown but have imputed values. The model treats known and unknown traits differently. The difficulty is that the known and unknown traits are unique for all species. For example, while some species might only be missing body mass data, others might miss other traits, or multiple other traits. The model, therefore, needs a lookup table that tells it which of a species’ trait are known and which are unknown.

```
#-- USE MEAN VALUES FOR KNOWN TRAITS
known.trait.values <- df.traits.mean.scaled[,v.trait.cov]
known.trait.values[!is.na(df.traits.sd[,v.trait.cov])] <- NA


#-- IDENTIFY IMPUTED TRAITS FOR EACH SPECIES
##-- Number of imputed traits per species
n.imputed.traits <- apply(known.trait.values, 1, function(x){sum(is.na(x))})


#-- Create Matrix of imputed trait indices
imputed.traits.index <-
  matrix(NA,
         ncol = max(n.imputed.traits),
         nrow = n.species,
         dimnames = list( "species" = v.species.list,
                          "index" = 1:max(n.imputed.traits)))
for (s in 1:n.species) {
  imputed.traits.index[s,1:n.imputed.traits[s]] <- which(is.na(known.trait.values[s, ]))
}


#-- IDENTIFY KNOWN TRAITS FOR EACH SPECIES
##-- Number of imputed traits per species
n.known.traits <- apply(known.trait.values, 1, function(x){sum(!is.na(x))})


#-- Create Matrix of known trait indices
known.traits.index <-
  matrix(NA,
         ncol = max(n.known.traits),
         nrow = n.species,
         dimnames = list( "species" = v.species.list,
                          "index" = 1:max(n.known.traits)))

for (s in 1:n.species) {
  known.traits.index[s,1:n.known.traits[s]] <- which(!is.na(known.trait.values[s, ]))
}
```

#### Species X Site lookup tables

The replicates used in the model are Species X Site combinations (e.g. a tapir living in site Asia\_1). Species can, thus, be repeated multiple times. In contrast, the trait data is a table where every species has exactly one row. This section creates a lookup table that tells the model which trait data corresponds with which Species X Sites data, to deal with these mismatched dimensions.

```
#---- CREATE ORDERED SEQUENCE OF ALL SPECIES IN THE TRAIT DATASET
#--------------------------------------------–
trait.order = data.frame(known.trait.values) # save as df
trait.order$Binomial = rownames(trait.order) # Add Rownames as a Column
trait.order$Index = sort(rownames(known.trait.values),
                         index.return = T)[[2]] # give the order of each species
trait.order = trait.order[,c("Binomial", "Index")] # save only relevant variables


#---- MAKE LIST OF PRESENCE DATASETS
#--------------------------------------------–

l.trait.index = list(df.pres.global[,c("Binomial", "Bio.Realm", "Present.Globally")],
                     df.pres.regional[,c("Binomial", "BRU", "Present.Regionally")],
                     df.pres.local[,c("Binomial", "Polygon.ID", "Present.Locally")])

names(l.trait.index) = scale.names # add name to list elements


#---- FILL LIST (AND MERGE IT WITH TRAIT DATA INDEX VALUES)
#--------------------------------------------–
for (i.scale in 1:length(scale.names)) {
  
  # select data frame
  df = l.trait.index[[i.scale]]
  
  # Merge Trait Index with Presence Data 
  trait.index = merge(df,
                      trait.order,
                      by = "Binomial",
                      all.x = T) # merge

  # Add Species Names as Rownames
  rownames(trait.index) = paste0(trait.index$Binomial,
                                 "_", trait.index[,2]) 
  
  # Reorder data so that species from the same site are next to each other (important!!!!)
  trait.index = trait.index[match(rownames(df),
                                  rownames(trait.index)),]
  
  # Save the Trait Index Separately (used later in model)
  temp = rownames(trait.index) # save rownames
  trait.index = as.vector(trait.index[,c("Index")]) # turn into vector
  names(trait.index) = temp # save rownames
  
  # Save df to list
  l.trait.index[[i.scale]] = trait.index
}
```

### Prepare the Model

#### Define Nimble Model

```
modelCode <- nimbleCode({
#-----------------------------------------------------------------------–
## – DEFINE PRIORS ––––
#-----------------------------------------------------------------------–  
  
#---- DEFINE PRIORS OF THE PSI-ESTIMATES 
#--------------------------------------------–
  
#~~~~~~ NOTE ~~~~~~~~~~~~~~~~~~~~~~~~~~~~~~~~~~~~~~~~~~~~~~~~~~~~~~~~~~~~~~~~~~~~
#
#   This section defines the priors for the psiRJ values. 
#   It creates a unique value for every scale. 
# 
#~~~~~~~~~~~~~~~~~~~~~~~~~~~~~~~~~~~~~~~~~~~~~~~~~~~~~~~~~~~~~~~~~~~~~~~~~~~~~~~~
      
  for(i.scale.data in 1:n.scales.data){
    psiRJ[i.scale.data] ~ dunif(0,1)
  } # close for loop (i.scale)
  
  
  
#---- DEFINE COEFFICIENT PRIORS
#----------------------------------–
  
#~~~~~~ NOTE ~~~~~~~~~~~~~~~~~~~~~~~~~~~~~~~~~~~~~~~~~~~~~~~~~~~~~~~~~~~~~~~~~~~~
#
#   This is where the priors for the model coefficients are specified. 
#   Note that every raw coefficient estimate is multiplied with a reversible
#   jump value (zRJ) that can take a value of 0 or 1.
#
#   If the value is 0, the coefficient has no influence on the model. 
#   If the value 1, the coefficient is included as normal. 
#
#   By estimating the zRJ values, the model can estimate which coefficients 
#   should be included in the model and which coefficients should be excluded.
# 
#~~~~~~~~~~~~~~~~~~~~~~~~~~~~~~~~~~~~~~~~~~~~~~~~~~~~~~~~~~~~~~~~~~~~~~~~~~~~~~~~
  
  ##---- Global Coefficients
  beta.global0  ~ dlogis(0, 1) # intercept
  
  for (i.cov in 1:n.trait.covs) { # define trait coefficients
    beta.global.traits.raw[i.cov] ~ dlogis(0, 1)
    zRJ.global.traits[i.cov] ~ dbern(psiRJ[1])
    beta.global.traits[i.cov] <- beta.global.traits.raw[i.cov] * zRJ.global.traits[i.cov]
  } # close for (i.cov)
  
  for (i.cov in 1:n.site.covs.global) { # define the other coefficients
    beta.global.site.raw[i.cov] ~ dlogis(0, 1)
    zRJ.global.site[i.cov] ~ dbern(psiRJ[2])
    beta.global.site[i.cov] <- beta.global.site.raw[i.cov] * zRJ.global.site[i.cov]
  } # close for (i.cov)
  
  
  ##---- Regional Coefficients
  beta.regional0  ~ dlogis(0, 1) # intercept
  
  for (i.cov in 1:n.trait.covs) { # define trait coefficients
    beta.regional.traits.raw[i.cov] ~ dlogis(0, 1)
    zRJ.regional.traits[i.cov] ~ dbern(psiRJ[3])
    beta.regional.traits[i.cov] <- beta.regional.traits.raw[i.cov] * zRJ.regional.traits[i.cov]
  } # close for (i.cov)
  
  for (i.cov in 1:n.site.covs.regional) { # define the other coefficients
    beta.regional.site.raw[i.cov] ~ dlogis(0, 1)
    zRJ.regional.site[i.cov] ~ dbern(psiRJ[4])
    beta.regional.site[i.cov] <- beta.regional.site.raw[i.cov] * zRJ.regional.site[i.cov]
  } # close for (i.cov)
  
  
  ##---- Local Coefficients
  beta.local0  ~ dlogis(0, 1) # intercept
  
  for (i.cov in 1:n.trait.covs) { # define trait coefficients
    beta.local.traits.raw[i.cov] ~ dlogis(0, 1)
    zRJ.local.traits[i.cov] ~ dbern(psiRJ[5])
    beta.local.traits[i.cov] <- beta.local.traits.raw[i.cov] * zRJ.local.traits[i.cov]
  } # close for (i.cov)
  
  for (i.cov in 1:n.site.covs.local) { # define the other coefficients
    beta.local.site.raw[i.cov] ~ dlogis(0, 1)
    zRJ.local.site[i.cov] ~ dbern(psiRJ[6])
    beta.local.site[i.cov] <- beta.local.site.raw[i.cov] * zRJ.local.site[i.cov]
  } # close for (i.cov)
  
  
  
#---- IMPUTE SPECIES TRAITS 
#----------------------------------–
  
#~~~~~~ NOTE ~~~~~~~~~~~~~~~~~~~~~~~~~~~~~~~~~~~~~~~~~~~~~~~~~~~~~~~~~~~~~~~~~~~~
#
#   This is where the uncertainty around the trait data is incorporated. 
#   Because parts of the trait data are uncertain, we are effectively treating them like priors. 
#   The model can sample their values from a distribution 
#   This section essentially creates the trait dataset the regressions will use,
#   with uniquely sampled values every iteration. 
#   It takes the known values and samples unknown values from the given distributions. 
#
#   Also note that the trait values are the same for all regressions,
#   so we do not do this separately for every scale.
# 
#~~~~~~~~~~~~~~~~~~~~~~~~~~~~~~~~~~~~~~~~~~~~~~~~~~~~~~~~~~~~~~~~~~~~~~~~~~~~~~~~
      
  
  for (i.sp in 1:n.species) { # loop through species
    #---- IMPUTED TRAIT VALUES
    for(i.cov in 1:n.imputed.traits[i.sp]){ # loop trough covs
      ar.trait.mean[i.sp, imputed.traits.index[i.sp,i.cov]] ~ dunif(-10,10)
      ar.trait.sd[i.sp, imputed.traits.index[i.sp,i.cov]] ~ dgamma(1,1)

      for (i.imp in 1:n.imputations) { # loop through samples
        imputed.trait.values[i.imp,i.sp,imputed.traits.index[i.sp,i.cov]] ~
          dnorm( mean = ar.trait.mean[i.sp,imputed.traits.index[i.sp,i.cov]],
                 sd = ar.trait.sd[i.sp,imputed.traits.index[i.sp,i.cov] ])
      } # close for loop (i.imp)

      cov.traits[i.sp,imputed.traits.index[i.sp,i.cov]] ~
        dnorm( mean = ar.trait.mean[i.sp, imputed.traits.index[i.sp,i.cov]],
               sd = ar.trait.sd[i.sp, imputed.traits.index[i.sp,i.cov]])
    } # close for loop (i.cov)

    #---- KNOWN TRAIT VALUES
    for(i.cov in 1:n.known.traits[i.sp]){ # loop trough covs
      cov.traits[i.sp,known.traits.index[i.sp,i.cov]] <-
        known.trait.values[i.sp, known.traits.index[i.sp,i.cov]]
    } # close for loop (i.cov)
  } #close for loop (i.sp)


  
  #-----------------------------------------------------------------------–
  ## – ESTIMATE SURVIVAL PROBABILITY ––––
  #-----------------------------------------------------------------------–  
  
#~~~~~~ NOTE ~~~~~~~~~~~~~~~~~~~~~~~~~~~~~~~~~~~~~~~~~~~~~~~~~~~~~~~~~~~~~~~~~~~~~~
#
#    This section estimates survival probability (psi).
# 
#~~~~~~~~~~~~~~~~~~~~~~~~~~~~~~~~~~~~~~~~~~~~~~~~~~~~~~~~~~~~~~~~~~~~~~~~~~~~~~~~~~
      
#----ESTIMATE GLOBAL SURVIVAL PROBABILITY
#------------------------------------------–
  
  for (i.ss in 1:n.global) {
    # Regression
    logit(psi.global[i.ss]) <- beta.global0 +
      inprod(beta.global.traits[1:n.trait.covs],
             cov.traits[trait.index.global[i.ss], 1:n.trait.covs]) +
      inprod(beta.global.site[1:n.site.covs.global],
             cov.sites.global[i.ss, 1:n.site.covs.global])
    
    # Estimate Presence
    isPresent.global[i.ss] ~ dbern(psi.global[i.ss])
  } # close for loop (i.ss)
  
  
#----ESTIMATE REGIONAL SURVIVAL PROBABILITY
#------------------------------------------–
  
  for (i.ss in 1:n.regional) {
    # Regression
    logit(psi.regional[i.ss]) <- beta.regional0 +
      inprod(beta.regional.traits[1:n.trait.covs],
             cov.traits[trait.index.regional[i.ss], 1:n.trait.covs]) +
      inprod(beta.regional.site[1:n.site.covs.regional],
             cov.sites.regional[i.ss, 1:n.site.covs.regional])
    
    # Estimate Presence
    isPresent.regional[i.ss] ~ dbern(psi.regional[i.ss])
    
  } # close for loop (i.ss)
  
  
#----ESTIMATE LOCAL SURVIVAL PROBABILITY
#------------------------------------------–
  
  for (i.ss in 1:n.local) {
    # Regression
    logit(psi.local[i.ss]) <- beta.local0 +
      inprod(beta.local.traits[1:n.trait.covs],
             cov.traits[trait.index.local[i.ss], 1:n.trait.covs]) +
      inprod(beta.local.site[1:n.site.covs.local],
             cov.sites.local[i.ss, 1:n.site.covs.local])
    
    # Estimate Presence
    isPresent.local[i.ss] ~ dbern(psi.local[i.ss])
    
  } # close for loop (i.ss)
  
})# close model definition
```

### Define Model Inputs

In the previous section we defined the structure of the model, but we have not provided any data to run the model. That is what we do here.

##### Define the Data

```
nimData <- list(
  
  # Presence Data
  isPresent.global = df.pres.global$Present.Globally,
  isPresent.regional = df.pres.regional$Present.Regionally, 
  isPresent.local = df.pres.local$Present.Locally,
  
  # Site Data
  cov.sites.global = as.matrix(l.site.data$Global),
  cov.sites.regional = as.matrix(l.site.data$Regional),
  cov.sites.local = as.matrix(l.site.data$Local),
  
  # Trait Data 
  imputed.trait.values = ar.traits.scaled[,,v.trait.cov]
)
```

##### Define the Constants

Some parameters in the model never change. They are considered constants rather than data. We define them here.

```
nimConstants <- list(
  
  n.species = n.species, # the number of species
  n.scales = length(scale.names), # the number of scales in the model
  n.scales.data = length(scale.names) * 2, # Nr of Scales X Nr of variable types (=2)
  
  # Replicates at each scale
  n.global = nrow(df.pres.global),
  n.regional = nrow(df.pres.regional),
  n.local = nrow(df.pres.local),
  
  # Traits
  n.trait.covs = length(v.trait.cov),
  n.imputations = dim(ar.traits.scaled)[1],
  n.imputed.traits = n.imputed.traits,
  n.known.traits = n.known.traits,
  
  imputed.traits.index = imputed.traits.index,
  known.traits.index = known.traits.index,
  known.trait.values = df.traits.mean.scaled[,v.trait.cov],
  
  trait.index.global = l.trait.index$Global,
  trait.index.regional = l.trait.index$Regional,
  trait.index.local = l.trait.index$Local,
  
  # Nr of Site (non-trait) Covariates at each scale
  n.site.covs.global = ncol(l.site.data$Global),
  n.site.covs.regional = ncol(l.site.data$Regional),
  n.site.covs.local = ncol(l.site.data$Local), 
  
  # Nr Of Sites for Each Scale
  n.sites.global = length(unique(df.sites$Bio.Realm)),
  n.sites.regional = length(unique(df.sites$BRU)),
  n.sites.local = length(unique(df.sites$Polygon.ID))
)
```

##### Define Initial Values

Here, we define the initial values of several of the model parameters (i.e. the starting values for the MCMC chains).

```
nimInits <- list(
  
  # Reversible Jump (take 1 as starting values)
  zRJ.global.traits = rep(1, nimConstants$n.trait.covs),
  zRJ.global.site = rep(1, nimConstants$n.site.covs.global),
  
  zRJ.regional.traits = rep(1, nimConstants$n.trait.covs),
  zRJ.regional.site = rep(1, nimConstants$n.site.covs.regional),
  
  zRJ.local.traits = rep(1, nimConstants$n.trait.covs),
  zRJ.local.site = rep(1, nimConstants$n.site.covs.local),
  
  # Coefficients (take 0 as starting values)
  beta.global0 = 0,
  beta.global.traits.raw = rep(0, nimConstants$n.trait.covs),
  beta.global.site.raw = rep(0, nimConstants$n.site.covs.global),
  beta.global.traits = rep(0, nimConstants$n.trait.covs),
  beta.global.site = rep(0, nimConstants$n.site.covs.global),
  
  beta.regional0 = 0,
  beta.regional.traits.raw = rep(0, nimConstants$n.trait.covs),
  beta.regional.site.raw = rep(0, nimConstants$n.site.covs.regional),      
  beta.regional.traits = rep(0, nimConstants$n.trait.covs),
  beta.regional.site = rep(0, nimConstants$n.site.covs.regional),
  
  beta.local0 = 0,
  beta.local.traits.raw = rep(0, nimConstants$n.trait.covs),
  beta.local.site.raw = rep(0, nimConstants$n.site.covs.local),
  beta.local.traits = rep(0, nimConstants$n.trait.covs),
  beta.local.site = rep(0, nimConstants$n.site.covs.local),
  
  # Trait Values (take means as starting values)
  ar.trait.mean = df.traits.mean.scaled[,v.trait.cov], 
  cov.traits = df.traits.mean.scaled[,v.trait.cov]
)
```

##### Specify the parameters to retrieve

Nimble does not automatically save all parameters in the model. Here, we specify which parameters the model should save for later use.

```
nimParams <- c(
  
  # Trait Variables
  "beta.global.traits",
  "beta.regional.traits",
  "beta.local.traits",
  
  "beta.global.traits.raw",
  "beta.regional.traits.raw",
  "beta.local.traits.raw",
  
  # Non-Trait Variables
  "beta.global.site",
  "beta.regional.site",
  "beta.local.site",
  
  "beta.global.site.raw",
  "beta.regional.site.raw",
  "beta.local.site.raw",
  
  # Reversible Jump Values
  "zRJ.global.traits",
  "zRJ.regional.traits",
  "zRJ.local.traits",
  
  "zRJ.global.site",
  "zRJ.regional.site",
  "zRJ.local.site",
  
  # Estimated Survival Probability
  "psi.global",
  "psi.regional",
  "psi.local",
  
  # Occurrence
  "isPresent.global",
  "isPresent.regional",
  "isPresent.local"
)
```

### Fit the Model

The model can take a while to run. So we will take keep track of the time it takes

```
ptm <- proc.time()
```

Next, we bring together the model structure, data, constants, and initial values.

```
model <- nimbleModel( code = modelCode,
                      constants = nimConstants,
                      data = nimData,
                      inits = nimInits,
                      check = F,    
                      calculate = F)
```

Next, we define the MCMC algorithm.

```
## Compile Model in C++
cmodel <- compileNimble(model)

## Configure MCMC algorithm
MCMCconf <- configureMCMC( model = model,
                           monitors = nimParams,
                           control = list(reflective = TRUE),
                           thin = df.model.specs$thin,
                           enableWAIC = TRUE)
```

The reversible jump needs its own separate MCMC algorithm, which we specify here.

```
configureRJ(MCMCconf,
            targetNodes = c('beta.global.traits.raw','beta.global.site.raw'),
            indicatorNodes = c("zRJ.global.traits", "zRJ.global.site"),
            control = list(mean = 0, scale = .2))

configureRJ(MCMCconf,
            targetNodes = c('beta.regional.traits.raw','beta.regional.site.raw'),
            indicatorNodes = c("zRJ.regional.traits", "zRJ.regional.site"),
            control = list(mean = 0, scale = .2))

configureRJ(MCMCconf,
            targetNodes = c('beta.local.traits.raw','beta.local.site.raw'),
            indicatorNodes = c("zRJ.local.traits", "zRJ.local.site"),
            control = list(mean = 0, scale = .2))

MCMC <- buildMCMC(MCMCconf)
```

The model needs to be compiled. Compiling the model means translating the “R model” into a “C++ model” which is computationally more efficient. Next, we load that “C++ model” into our working environment as a distinct object that can be handled in R.

```
cMCMC <- compileNimble( MCMC,
                        project = model,
                        resetFunctions = TRUE)
```

Now, we are finally ready to fit the model.

```
myNimbleOutput <- runMCMC( mcmc = cMCMC,
                           nburnin = df.model.specs$nburnin,
                           niter = df.model.specs$niter,
                           nchains = df.model.specs$nchains,
                           WAIC = F,
                           inits = nimInits,
                           samplesAsCodaMCMC = TRUE)
```

```
Running chain 1 ...
```

```
|-------------|-------------|-------------|-------------|
|-------------------------------------------------------|
```

```
Running chain 2 ...
```

```
|-------------|-------------|-------------|-------------|
|-------------------------------------------------------|
```

```
Running chain 3 ...
```

```
|-------------|-------------|-------------|-------------|
|-------------------------------------------------------|
```

Finally, let’s check how long it took to run the model.

```
TotalRuntime <- proc.time()-ptm
print(TotalRuntime)
```

```
   user  system elapsed 
301.886   3.010 306.224
```

### Process Nimble Output

We need to process the Nimble output, because it is a bit unwieldy at the moment. Here, we use a custom function (provided with the data objects) to process the Nimble MCMC output into an object that can be more easily queried.

```
source("./Vignette_Submission_files/data/ProcessCodaOutput.R")
myNimbleOutputCoda <- ProcessCodaOutput(myNimbleOutput, DIC = F)
```

To get hold of parameters type: `myNimbleOutputCoda$sims.list$...`

#### Show Traceplots

Select which parameters to show trace plot of (this become really slow if we plot all PSI values, so we need to be selective). For visualisation purposes, we only show the first three plots.

```
## Select parameters to plot (here, all coefficients)
v.params = c(grep("beta.global", colnames(myNimbleOutput[[1]]), value = T),
             grep("beta.regional", colnames(myNimbleOutput[[1]]), value = T),
             grep("beta.local", colnames(myNimbleOutput[[1]]), value = T))

## Plot Trace Plots 
plot(myNimbleOutput[ ,v.params[1:3]]) # remove indexing to show all plots
```

###### PLOT TRACE PLOTS BUT WITH 0’S FROM REVERSIBLE JUMP REMOVED

Interpreting the traceplots can be difficult when using the reversible jump, because when parameters are not included, their value switches back to 0. This create messy trace plots. To provide plots that are easier to interpret, it can be helpful to plot traceplots where the values are excluded if the reversible jump excluded the parameter.

```
##– GLOBAL
zRJ.global <- myNimbleOutputCoda$sims.list$zRJ.global.traits # extract ZRJ trait data
colnames(zRJ.global) <- v.trait.cov # give colnames

df.cov.est.global = myNimbleOutputCoda$sims.list$beta.global.traits # get the data
colnames(df.cov.est.global) = dimnames(nimData$imputed.trait.values)[[3]] # give colnames
temp = zRJ.global[, dimnames(nimData$imputed.trait.values)[[3]]] # select relevant variables
df.cov.est.global[temp == 0] = NA # remove 0s
df.cov.est.global = data.frame(df.cov.est.global) # turn into DF
df.cov.est.global$Iteration = rep(1:nrow(myNimbleOutput$chain1), length(myNimbleOutput))
df.cov.est.global$Chain = as.factor(rep(1:length(myNimbleOutput), each = nrow(myNimbleOutput$chain1)))


##– REGIONAL
zRJ.regional<-myNimbleOutputCoda$sims.list$zRJ.regional.traits # extract ZRJ trait data
colnames(zRJ.regional)<- v.trait.cov # give colnames

df.cov.est.regional = myNimbleOutputCoda$sims.list$beta.regional.traits # get the data
colnames(df.cov.est.regional) = dimnames(nimData$imputed.trait.values)[[3]] # give colnames
temp = zRJ.regional[, dimnames(nimData$imputed.trait.values)[[3]]] # select relevant variables
df.cov.est.regional[temp == 0] = NA # remove 0s
df.cov.est.regional = data.frame(df.cov.est.regional) # turn into DF
df.cov.est.regional$Iteration = rep(1:nrow(myNimbleOutput$chain1), length(myNimbleOutput))
df.cov.est.regional$Chain = as.factor(rep(1:length(myNimbleOutput), each = nrow(myNimbleOutput$chain1)))


##– LOCAL
zRJ.local<-myNimbleOutputCoda$sims.list$zRJ.local.traits # extract ZRJ trait data
colnames(zRJ.local)<- v.trait.cov # give colnames

df.cov.est.local = myNimbleOutputCoda$sims.list$beta.local.traits
colnames(df.cov.est.local) = dimnames(nimData$imputed.trait.values)[[3]]
temp = zRJ.local[, dimnames(nimData$imputed.trait.values)[[3]]] # select relevant variables
df.cov.est.local[temp == 0] = NA
df.cov.est.local = data.frame(df.cov.est.local) # turn into DF
df.cov.est.local$Iteration = rep(1:nrow(myNimbleOutput$chain1), length(myNimbleOutput))
df.cov.est.local$Chain = as.factor(rep(1:length(myNimbleOutput), each = nrow(myNimbleOutput$chain1)))


## Prepare Colours
n <- df.model.specs$nchains
qual_col_pals = brewer.pal.info[brewer.pal.info$category == 'qual',]
col_vector = unlist(mapply(brewer.pal, qual_col_pals$maxcolors, rownames(qual_col_pals)))


##- PLOT GLOBAL
plot.list.global <- vector("list", length = ncol(df.cov.est.global)) # create list
for (i.var in 1:(ncol(df.cov.est.global)-2)){
  # Select Variable
  col.name = colnames(df.cov.est.global)[i.var] # define column that needs to be selected
  # Plot
  p = ggplot(df.cov.est.global) +
    geom_line(aes(x = Iteration, y = !!sym(col.name), col = Chain), linewidth = 0.35) +
    scale_color_manual(values=col_vector)
  # Save Output
  plot.list.global[[i.var]] = p
} # close loop


##- PLOT REGIONAL
plot.list.regional <- vector("list", length = ncol(df.cov.est.regional)) # create list
for (i.var in 1:(ncol(df.cov.est.regional)-2)){
  # Select Variable
  col.name = colnames(df.cov.est.regional)[i.var] # define column that needs to be selected
  # Plot
  p = ggplot(df.cov.est.regional) +
    geom_line(aes(x = Iteration, y = !!sym(col.name), col = Chain), linewidth = 0.35) +
    scale_color_manual(values = col_vector)
  # Save Output
  plot.list.regional[[i.var]] = p
} # close loop


##- PLOT LOCAL
plot.list.local <- vector("list", length = ncol(df.cov.est.local)) # create list
for (i.var in 1:(ncol(df.cov.est.local)-2)){
  # Select Variable
  col.name = colnames(df.cov.est.local)[i.var] # define column that needs to be selected
  # Plot
  p = ggplot(df.cov.est.local) +
    geom_line(aes(x = Iteration, y = !!sym(col.name), col = Chain), linewidth = 0.35) +
    scale_color_manual(values= col_vector)
  # Print Output
  plot.list.local[[i.var]] = p
} # close loop

plotlist.total = c(plot.list.global, plot.list.regional, plot.list.local)
```

We will plot the first 3 plots.

```
plotlist.total[1:3]
```

```
[[1]]
```

```
[[2]]
```

```
[[3]]
```

#### Plot Coefficients

First, we extract the posterior estimates of the coefficients for the global, regional and local scale.

```
l.coefficients = vector(mode = "list", length = length(scale.names))
names(l.coefficients) = scale.names

##- GLOBAL
betas.global <- myNimbleOutputCoda$sims.list[grepl("beta.global",
                                                   names(myNimbleOutputCoda$sims.list))]
betas.global = betas.global[!grepl(".raw",
                                   names(betas.global))] # remove .raw betas

colnames(betas.global[[1]]) <-colnames(l.site.data$Global)
colnames(betas.global[[2]]) <- v.trait.cov
betas.global = do.call(cbind, betas.global)

zRJ.global<-myNimbleOutputCoda$sims.list[grepl("zRJ.global", names(myNimbleOutputCoda$sims.list))]
colnames(zRJ.global[[1]]) <- colnames(l.site.data$Global)
colnames(zRJ.global[[2]]) <- v.trait.cov
zRJ.global = do.call(cbind, zRJ.global)
betas.global[zRJ.global==0]<-NA # turn coefficient values that were exlcuded by the reversible jump to NA
l.coefficients[["Global"]] = betas.global # save in list


##- REGIONAL
betas.regional <- myNimbleOutputCoda$sims.list[grepl("beta.regional", names(myNimbleOutputCoda$sims.list))]
betas.regional = betas.regional[!grepl(".raw", names(betas.regional))] # remove .raw betas
colnames(betas.regional[[1]]) <- colnames(l.site.data$Regional)
colnames(betas.regional[[2]]) <- v.trait.cov
betas.regional = do.call(cbind, betas.regional)

zRJ.regional<-myNimbleOutputCoda$sims.list[grepl("zRJ.regional", names(myNimbleOutputCoda$sims.list))]
colnames(zRJ.regional[[1]]) <- colnames(l.site.data$Regional)
colnames(zRJ.regional[[2]]) <- v.trait.cov
zRJ.regional = do.call(cbind, zRJ.regional)
betas.regional[zRJ.regional==0]<-NA # turn coefficient values that were exlcuded by the reversible jump to NA
l.coefficients[["Regional"]] = betas.regional # save in list


##- LOCAL
betas.local <- myNimbleOutputCoda$sims.list[grepl("beta.local", names(myNimbleOutputCoda$sims.list))]
betas.local = betas.local[!grepl(".raw", names(betas.local))] # remove .raw betas
colnames(betas.local[[1]]) <-  colnames(l.site.data$Local)
colnames(betas.local[[2]]) <- v.trait.cov
betas.local = do.call(cbind, betas.local)

zRJ.local<-myNimbleOutputCoda$sims.list[grepl("zRJ.local", names(myNimbleOutputCoda$sims.list))]
colnames(zRJ.local[[1]]) <-  colnames(l.site.data$Local)
colnames(zRJ.local[[2]]) <- v.trait.cov
zRJ.local = do.call(cbind, zRJ.local)
betas.local[zRJ.local==0]<-NA # turn coefficient values that were exlcuded by the reversible jump to NA
l.coefficients[["Local"]] = betas.local # save in list
```

Next, we want to plot these estimates. We create a file contains all the above data in a format suitable for ggplot. We also assign each coefficient to a category (Biogeographical, Trait, Local, Phylogeny), and some other data which will be helpful for plotting.

```
##- CREATE EMPTY VECTOR
l.coeff = vector(mode = "list", length = length(scale.names))
names(l.coeff) = scale.names

for(i.scale in 1:length(scale.names)){  
  
  #- SELECT BETAS & CONVERT INTO APPROPRIATE FORMAT
  betas = l.coefficients[[i.scale]]
  
  betas.df<-melt(betas)
  names(betas.df)<-c("idx","variable","beta")
  
  betas.aggr <- betas.df %>% 
    group_by(variable) %>% 
    summarise(p.inclusion= mean(!is.na(beta)))
  
  betas.df<-merge(betas.df,betas.aggr)
  
  ## ADD DIRECTIONS (create lookup table that will later be merged)
  df.direction = data.frame(colnames(betas)) # turn into df
  df.direction$Direction = "Negative" # add direction variable
  df.direction[which(apply(betas, 2, mean, na.rm = T) > 0), "Direction"] = "Positive" # add direction variable
  
  ## Add scale variable
  betas.df$scale = scale.names[i.scale]
  betas.df = merge(betas.df, df.direction, by.x = "variable", by.y = "colnames.betas.", all.x = T)
  
  
  ## Add Variable Type
  betas.df[, "variable.type"] = "Biogeographical"
  
  m = which(grepl(betas.df$variable, pattern = paste(v.trait.cov,collapse="|"))) ## Intrinsic Variables
  betas.df[m, "variable.type"] = "Traits"
  
  m = which(grepl(betas.df$variable, pattern = "Dim")) ## Intrinsic Variables
  betas.df[m, "variable.type"] = "Phylogeny"
  
  m = which(betas.df$variable %in% as.vector(outer(v.trait.cov, v.trait.cov, paste, sep=":"))) ## Intrinsic Variables
  betas.df[m, "variable.type"] = "Traits"
  
  m = which(grepl(betas.df$variable, pattern = paste(v.ext.cov.local,collapse="|"))) ## environmental variables
  betas.df[m, "variable.type"] = "Local"
  
  betas.aggr <- betas.df %>%
    group_by(variable) %>%
    summarise(p.inclusion = mean(!is.na(beta)))
  
  betas.df <- merge(betas.df,betas.aggr)
  
  l.coeff[[i.scale]] = betas.df
  
} # close for loop (i.scale)
```

We turn the list elements into a data frame

```
## Turn all list elements into df
df.betas = do.call(rbind, l.coeff) 

## Turn scale into factor and define order
df.betas$scale = factor(df.betas$scale, levels = scale.names)
df.betas$variable.type = factor(df.betas$variable.type,
                                levels = c("Biogeographical",
                                           "Traits",
                                           "Phylogeny",
                                           "Local")) 

## Set The levels in the scale column
levels(df.betas$scale) = scale.names
```

We change the names of variable so that they are easier to interpret in the plot.

```
##- Change names of variables for plotting
df.betas$variable = as.character(df.betas$variable)

df.betas[which(grepl("Brain.Size", df.betas$variable)), "variable"] = "Brain Volume"
df.betas[which(grepl("Generation.Length", df.betas$variable)), "variable"] = "Generation Length"
df.betas[which(grepl("Mass.kg.log", df.betas$variable)), "variable"] = "Body Mass"
df.betas[which(grepl("Stratum.Numeric", df.betas$variable)), "variable"] = "Scancoriality"
df.betas[which(grepl("Diet.Vertebrate", df.betas$variable)), "variable"] = "Carnivory"
df.betas[which(grepl("Population.Density", df.betas$variable)), "variable"] = "Population Density"
df.betas[which(grepl("Forest.Cover", df.betas$variable)), "variable"] = "Forest Cover"
df.betas[which(grepl("WDPA.", df.betas$variable)), "variable"] = "Coverage by PA"
df.betas[which(grepl("Africa", df.betas$variable)), "variable"] = "Africa"
df.betas[which(grepl("Dim1", df.betas$variable)), "variable"] = "Phylo. Dim1"
df.betas[which(grepl("Dim2", df.betas$variable)), "variable"] = "Phylo. Dim2"
```

We rename the names of the scales (global, regional, local) so that they also include the sample size of each regression.

```
##- ADD SCALE NAME WITH SAMPLE SIZE 
df.betas[which(df.betas$scale == "Global"), "scale.n.size"] =
  paste0("Global (n = ", nrow(df.pres.global),")")

df.betas[which(df.betas$scale == "Regional"), "scale.n.size"] =
  paste0("Regional (n = ", nrow(df.pres.regional),")")

df.betas[which(df.betas$scale == "Local"), "scale.n.size"] =
  paste0("Local (n = ", nrow(df.pres.local),")")


df.betas$scale.n.size = factor(df.betas$scale.n.size,
                           levels  = c(paste0("Global (n = ", nrow(df.pres.global),")"),
                                       paste0("Regional (n = ", nrow(df.pres.regional),")"),
                                       paste0("Local (n = ", nrow(df.pres.local),")")))
```

We turn the variable names into variables and order them for the plotting.

```
temp = unique(df.betas[,c("variable.type", "variable")])
df.betas$variable = factor(df.betas$variable,
                           levels = temp[order(temp$variable.type, temp$variable),"variable"])
```

We want to highlight those variables that are included more than 50% of the time. So we create a variable to identify them.

```
##- Turn inclusion into percentages
df.betas$p.inclusion = df.betas$p.inclusion * 100

beta.sel <- df.betas %>% 
  group_by(variable) %>% 
  filter(p.inclusion > 50)
```

and now we finally create the plot.

```
##- Use the quantiles of the coefficient estimates to determine x-limits
x.lim= quantile(abs(df.betas$beta), probs = c(0.999), na.rm = T)

##- Plot
(p.coeff = ggplot() +
    geom_violin(data = df.betas,
                aes(beta, variable, fill= p.inclusion),
                draw_quantiles = c(0.025, 0.5, 0.975),
                lwd = 0.2,
                col = "darkgrey",
                scale = "width") +
    geom_violin(data = beta.sel,
                aes(beta, variable, fill= p.inclusion),
                draw_quantiles = c(0.025, 0.5, 0.975),
                lwd = 0.4,
                col = "black",
                scale = "width") +
    geom_vline(xintercept = 0,
               lty = "dashed",
               col = "black",
               lwd = 0.4) +
    xlab(NULL) +
    ylab(NULL) +
    labs(fill = "Iterations Included (%)") +
    xlim(c(-x.lim, x.lim)) +
    theme(legend.position= "bottom",
          legend.title = element_text(vjust = .8),
          legend.title.align=-300,
          panel.grid.major = element_line(color = 'grey87',
                                          linetype = 'solid',
                                          size = 0.3),
          panel.grid.minor = element_blank(),
          strip.text.x = element_text(size=12, face="bold"),
          strip.text.y = element_text(size=8),
          strip.background = element_rect(colour="black", fill="#d8e6c5"),
          panel.background = element_rect(fill = "white",
                                          colour = "grey1"),
          legend.box.background = element_blank()) +
    scale_fill_gradient(low = "white",
                        high = "#516b2d") +
    facet_grid( variable.type ~ scale.n.size,
                drop = T,
                scales = "free_y",
                space = "free"))
```
