## Supplementary figures and images for "Predictors of extinction risk in large tropical forest mammals: from global to local"

### SI_Data_1_rarefaction_curves.pdf

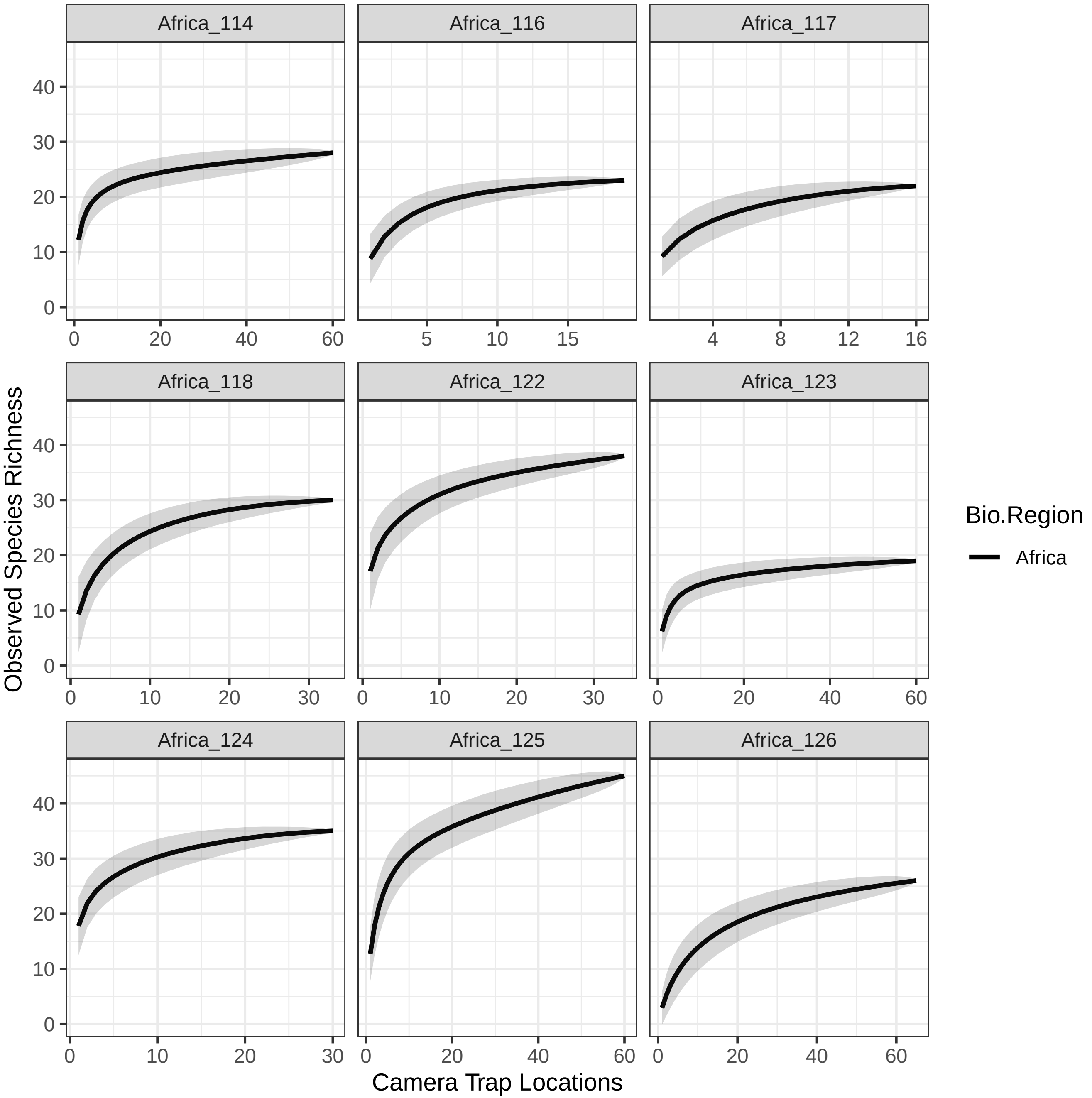

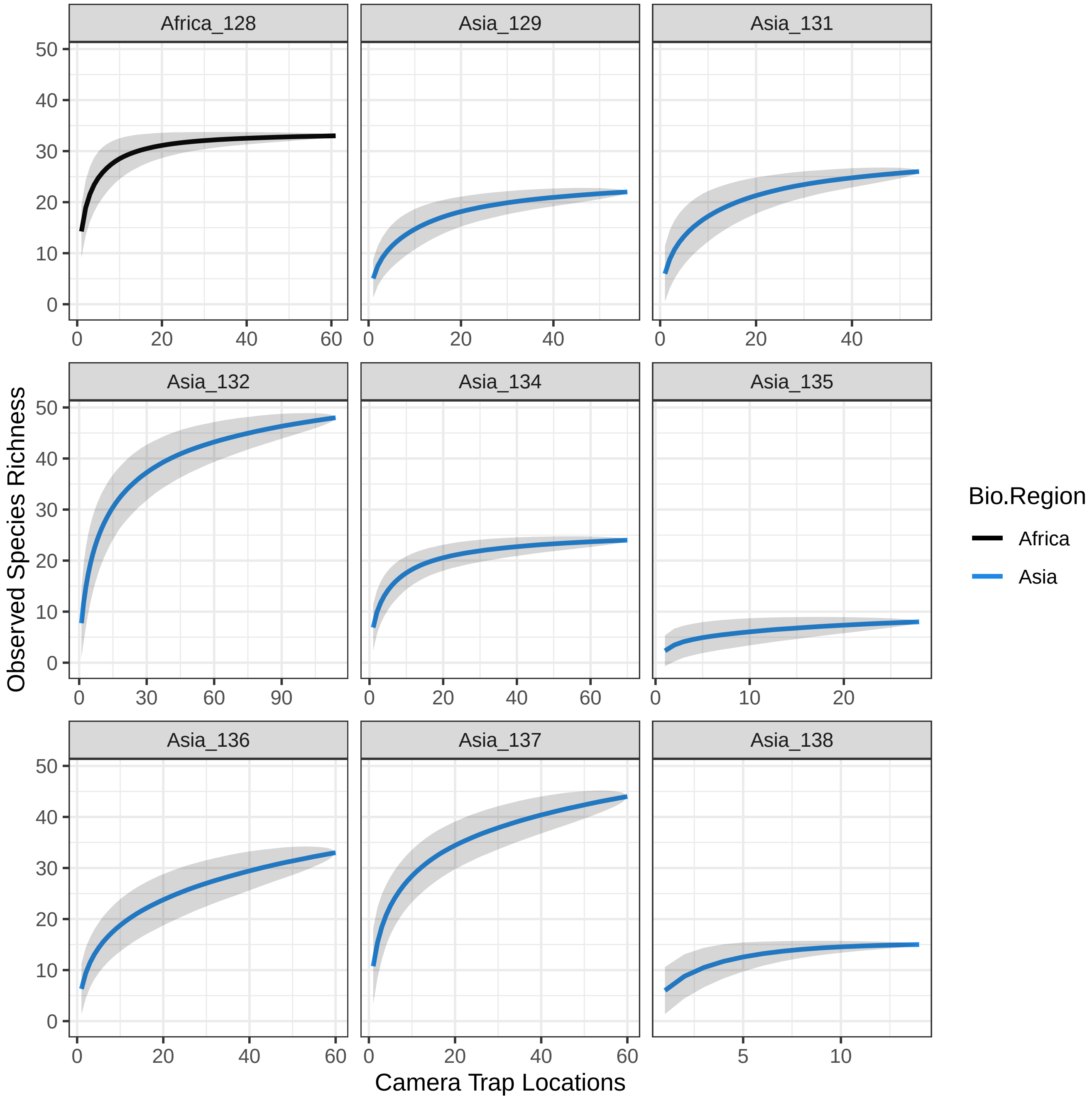

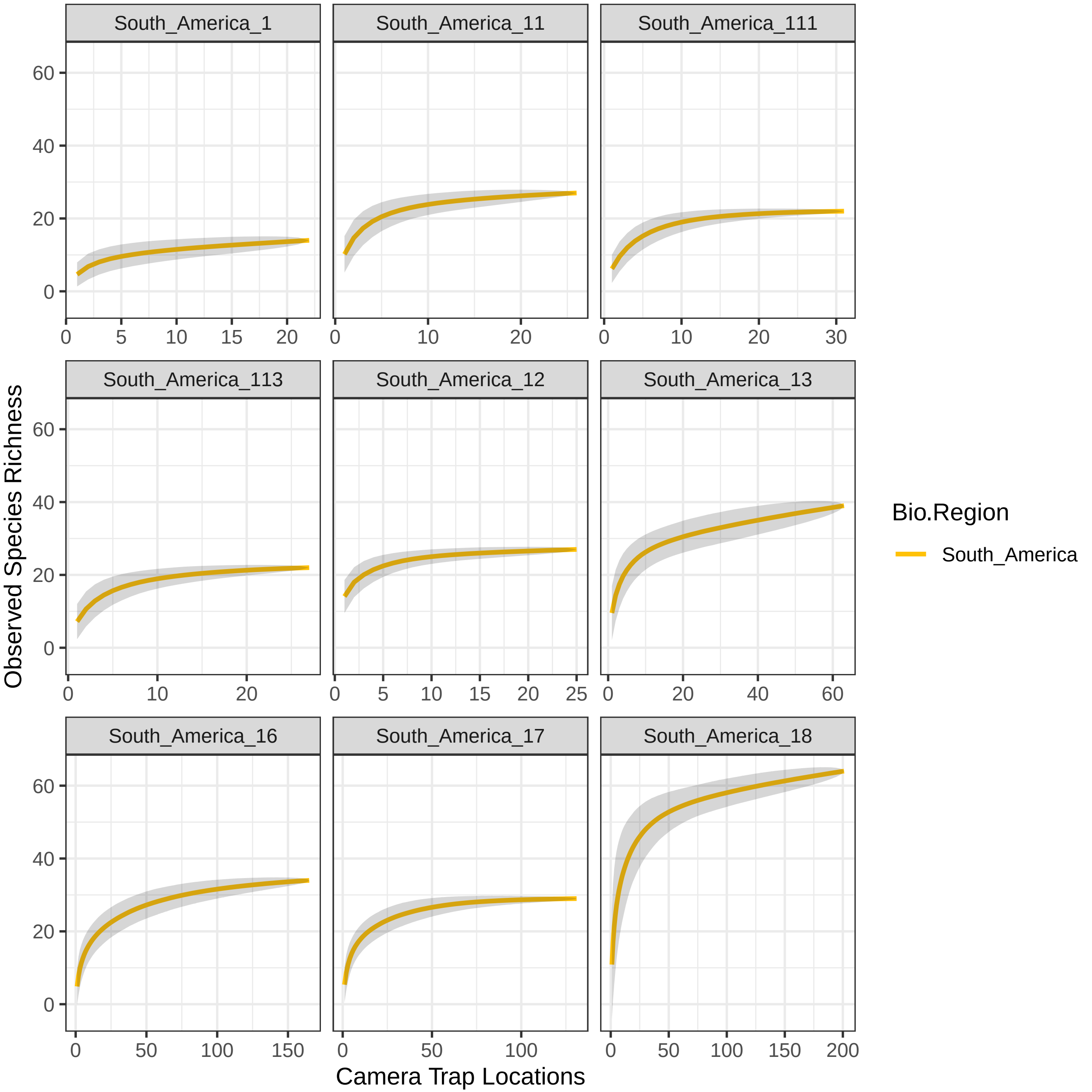

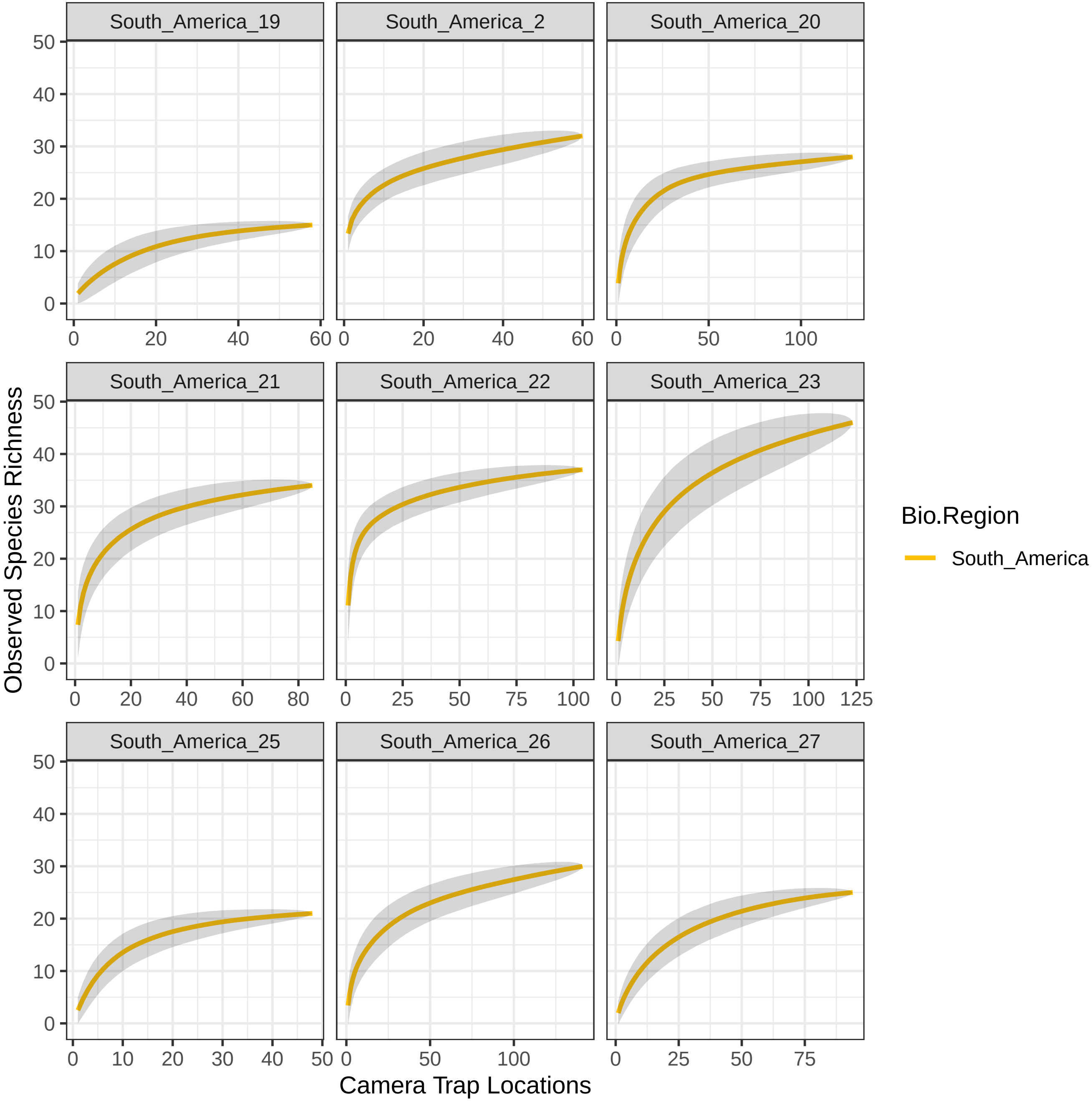

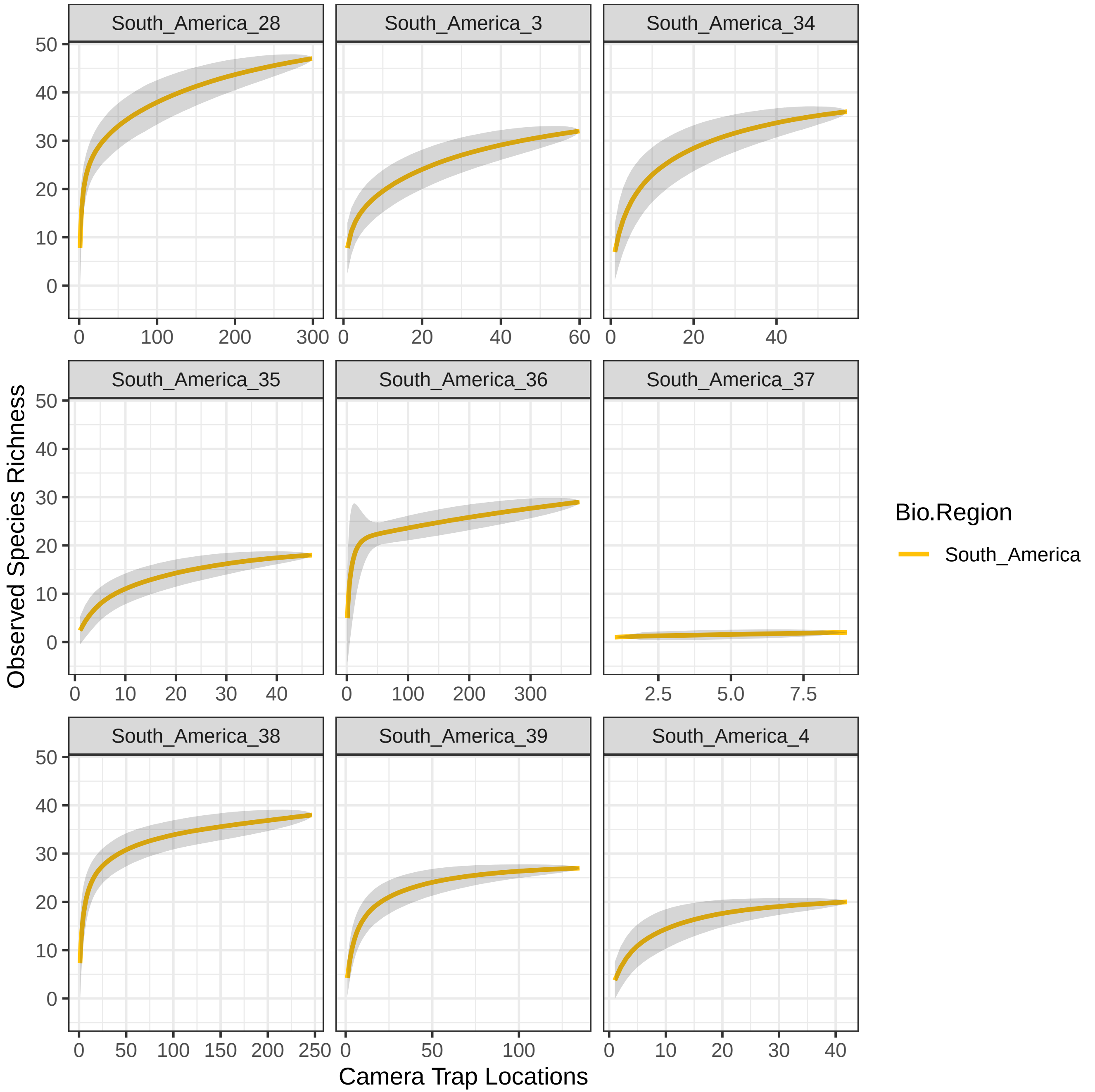

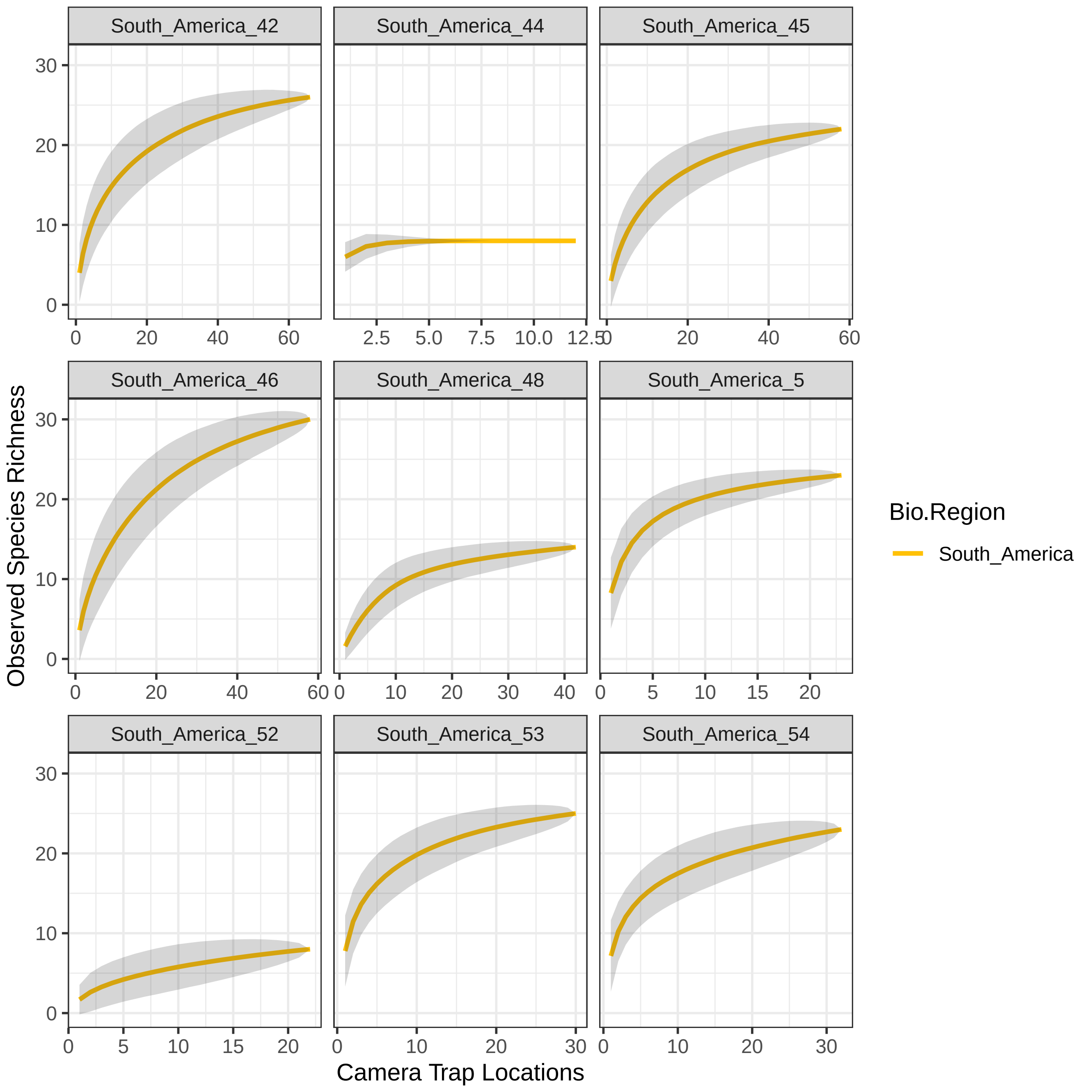

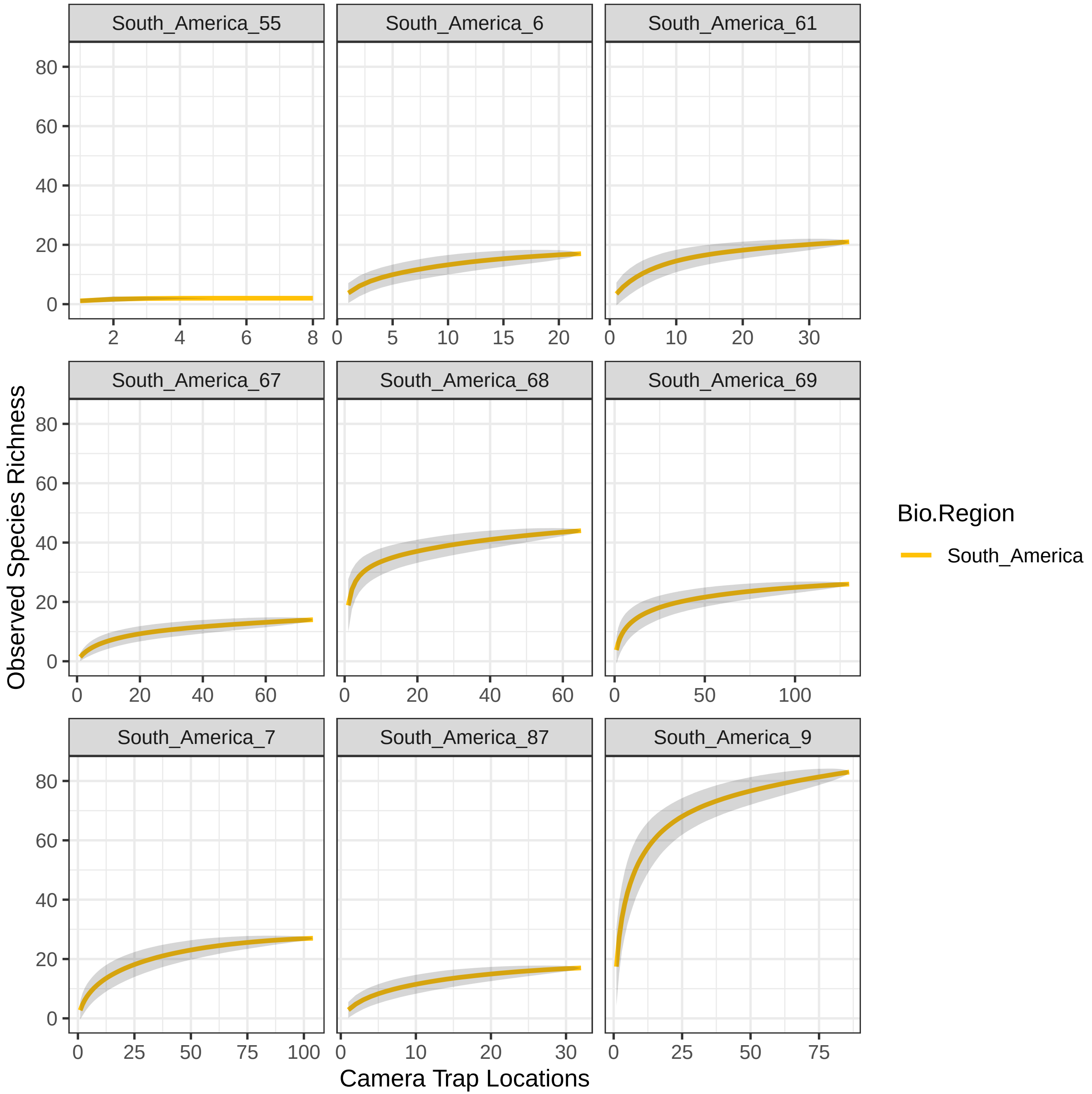
